# Dynamic cholesterol redistribution favors membrane fusion pore constriction

**DOI:** 10.1101/2022.04.15.488512

**Authors:** Andrew H. Beaven, Kayla Sapp, Alexander J. Sodt

## Abstract

Previous experiments have shown that cholesterol strongly prefers concave leaflets (which have negative curvature and are typically thin), but cholesterol also orders and thickens bilayers (promoting liquid-ordered phases with positive curvature). Our all-atom molecular dynamics simulations resolve this discrepancy for highly curved fusion pores, similar to those found in the nascent fusion and terminal fission steps of endo-/exocytosis. We find that cholesterol is strongly excluded by bilayer thinning in the fusion pore neck, which is caused by the neck’s net negative Gaussian (saddle) curvature. Consistent with experiment and our fusion pore simulations, analysis of liquid-disordered planar bilayers indicates that cholesterol prefers overall thicker *bilayers*, but negative *leaflet* curvature. The exclusion of cholesterol from the neck because of saddle Gaussian curvature implies that it helps drive fusion pore closure, consistent with literature evidence that membrane reshaping is connected to lateral phase separation.

## I. INTRODUCTION

Membranes in both endo- and exocytosis proceed through a highly strained *fusion pore* [1–3] stage, regardless of the protein machinery that assists the specific process. These pore structures arise, for example, during vesicle fusion to release neurotransmitters [4, 5] and preceding the fission step of clathrin-mediated endocytosis [6, 7]. It is predicted that formation of intermediates along the fusion/fission pathway have energetic barriers requiring tens to hundreds of *k*_B_*T* to surpass [8–11]. Nature uses a diverse set of proteins, performing specific tasks, to help overcome these large barriers during endo-/exocytosis processes: clathrin and adaptor proteins for forming buds with fusion pore necks [7]; dynamin to complete fission of the budded vesicle [12]; ESCRT-III for outward vesicular budding from cells [13]; COPII for vesicular budding from the endoplasmic reticulum, allowing transport to the Golgi apparatus [14]; and viral proteins that initiate enveloped virus fusion with the cell, such as SARS-CoV-2 spike proteins [15]. Major questions persist regarding how protein and lipid components stabilize strained intermediates. Why does cholesterol (chol), which stiffens membranes, support endocytosis [16]? What is the biophysical basis of sterol perturbations on viral entry and budding [17]? Can membrane mechanics be perturbed to favor the rapid opening and closing of pores, such as in “kiss-and-run” endocytosis [5]?

Energetic models, supported by experimental validation, predict the pathway for pore formation and closure [2, 18, 19]. The standard model for quantifying membrane shape energy is the Helfrich/Canham (HC) Hamiltonian [20, 21], a spring-equivalent model of the deformation, which is quantified by the total curvature (*J* = *c*_1_ + *c*_2_) and Gaussian curvature (*K* = *c*_1_*c*_2_). Here *c*_1_ and *c*_2_ are the orthogonal principal curvatures of the surface, describing parabolic deviations away from the plane. Lipid chemical identity favors specific curvatures, quantified in the spontaneous curvature (*J*_0_) [22–24]. The bending modulus (*κ*) determines the stiffness of the membrane, setting the overall energy scale [25–27]. The Gaussian curvature modulus (*κ*_G_) determines the coupling between principal curvature directions, differentiating, for example, spheres from tubes that might otherwise have the same *J* [8, 21, 28–30]. The model energy for a bilayer integrated at the midplane surface (*S*) is:

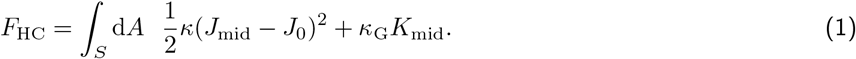

As an example, a small cylinder (e.g., a nascent fusion pore neck) with a height of 20 Å, a radius of 20 Å (*J* = (20 Å)^*−*1^, *K* = 0), and a bending modulus of 20 kcal/mol, yields an *F*_HC_ *≈* 63 kcal/mol, which is attainable by dynamin and ESCRTIII. Moving from the cylindrical neck and into the pore’s hourglass shape, pore structure energetics are strongly determined by *κ*_G_. It is hypothesized that particular lipid mixtures can bring *κ*_G_ to zero [8, 31], making the lipids potent “fusogens.” Thus, enrichment/depletion of lipids that influence *J*_0_ and *κ*_G_ (e.g., possible fusogens), for example, can dramatically reduce endo-/exocytosis barriers.

This work focuses on chol. A body of literature has demonstrated that chol chemistry and concentration dramatically affect virus-endosome fusion [17, 32], as well as plasma membrane fusion and fission [16, 33]. Understanding chol’s role in mechanics is complicated by disparate observations. Chol prefers negative *J*_0_ [34], implying that it lowers the barrier to fusion by enriching in the fusion pore. However, chol prefers ordered, stiff bilayers, implying that it would be depleted from highly bent fusion pores. It is possible that with its dual roles as ordering agent and negative *J*_0_ lipid, chol may serve different purposes at different stages of reshaping. Indeed, its ability to rapidly flip between the leaflets of membranes makes it uniquely suited to rapidly relieve stresses [35–37].

It is an open question to what extent *J*_0_ and *κ*_G_ are determined by individual or collective lipid properties – a stark case being lateral lipid phase separation. It is currently hypothesized that cellular outer leaflets exist near a liquid-ordered/liquid-disordered (L_o_/L_d_) transition because of the leaflets’ high saturated tail and chol content [38–40]. In an additive model [41], single lipid properties are compositionally weighted to determine an entire region’s mechanical properties. In reality, L_o_ and L_d_ phase mechanical properties are dramatically different [42], and individual lipids have “local extents” to which they influence their surroundings [41, 43]. In any system, an L_o_/L_d_ phase separation creates a line tension because of hydrophobic mismatch between the thicker L_o_ and thinner L_d_. Phase separation, line tension, and curvature are coupled [35, 42, 44–46] and can result in pore closure [16, 33, 47, 48]; chol is central to each of these elements. The simulations in this work link the geometric thinning of the hydrophobic interior to a driving force for phase separation (the difference in thickness between phases).

We simulated two pores to infer the thermodynamic effect of lipid composition, in particular chol, on pore stability. Previous simulations of fusion pores and intermediates have typically focused on structural dynamics and intermediate evolution, and therefore, have necessarily used coarse-grained resolution and/or simple compositions [49–56]. Herein, we focus on the energetics of a metastable, nascent fusion pore structure, similar to the work by Cui & coworkers [57]. In that work, the authors established the correspondence of molecular simulations and continuum models at very small lengthscales, and in one case observed substantial thinning at the fusion pore neck. Instead, herein, simulations were performed at all-atom resolution with a multi-component, asymmetric, plasma membrane mimetic. The pores simulated here change shape considerably slower than the relaxation timescale of composition in the neck (*∼*200 ns; see the Results section on timescales), which allows the simulation to quantify enrichment of lipids on structures whose shape is out of equilibrium. Fusion pore shape relaxation requires relaxation of both water across the bilayer (osmotic and hydrostatic stress), as well as relaxation of lipids in the inner and outer leaflet (differential stress [37]). While we report the slowly changing shape of the fusion pores here, this work focuses on the redistribution of lipids on nearly static pores.

Independent, continuous *∼*5.5 µs simulations of multi-component, asymmetric pores simulated on Anton 2 [58] demonstrate that chol is excluded from both leaflets of the fusion pore’s neck – an unexpected result for a lipid with strong negative *J*_0_. We show that chol exclusion is driven primarily by its preference for a bilayer with a thicker interior and that the neck bilayer is *thinned* relative to the bulk. We further demonstrate that rather than favoring reshaping broadly, chol drives fusion pore *collapse*, that is, smaller fusion pores, consistent with early theory on *κ*_G_ [44, 45, 59]. Thus, chol accumulation should favor endocytosis (e.g., viral entry through the plasma membrane) and budding [16, 33, 34, 44–46, 60], but disfavor viral escape from the endosome that requires expansion to a wide fusion pore [17, 61, 62]. However, unlike previous models [16, 47, 48] that proposed fission would occur because of laterally phase separated domains, our model only assumes that: 1) chol prefers thicker bilayers, and 2) fusion pore necks are thinned according to a straightforward mathematical and mechanical analysis.

The evidence and explanation of anomalous chol redistribution in fusion pores is presented as follows: First, geometric relations explain how curvature at the leaflet neutral surface (*δ*) and bilayer hydrophobic thickness (*t*) depend on shape at the bilayer midplane. Next, methodology is shown for how a literature-informed HC model is applied to treat expected redistribution. The lipid distributions in the fusion pore necks determined by molecular dynamics simulations are compared to the literature-informed model, highlighting the unexpected observed chol depletion in the pore neck. We then propose that hydrophobic thinning in the pore neck due to saddle Gaussian curvature is the mechanical basis for chol depletion. Finally, theory is presented for how dynamic redistribution of chol acts to shrink pore structures, favoring the late stages of vesicle budding.

## II. METHODS

### A. Build and simulation

The multi-component, planar bilayer with asymmetric leaflets was built using scripts from CHARMM-GUI [63]. We define the outer leaflet as the exoplasmic leaflet that contains chol (30%), monosialoganglioside (18:0/sphingosine; GM3 = 10%), palmitoyl-sphingomyelin (16:0/sphingosine; PSM = 28%), palmitoyl-arachidonoyl-phosphatidylcholine (16:0/20:4; PAPC = 7%), and palmitoyl-linoleoyl-phosphatidylcholine (16:0/18:2; PLPC = 25%). The inner leaflet is defined as the cytoplasmic leaflet that contains chol (30%), PLPC (16%), stearoyl-arachidonoyl-phosphatidylethanolamine (18:0/20:4; SAPE = 34%), and stearoyl-arachidonoyl-phosphatidylserine (18:0/20:4; SAPS = 20%). The system was neutralized and brought to *∼*150 mM KCl concentration. Starting from two *F*_HC_ minimized continuum mesh surfaces (one with a radial constraint), asymmetric fusion pores with different radii (P_small_ and P_large_) were built using in-house code that bends and grafts pieces of the planar bilayer onto the curved fusion pore geometry. Using this methodology, the bilayer and two pores had approximately the same lipid compositions and solution salt concentration. Tables S1–S3 of the Supplemental Material contain lipid concentrations for each system and each leaflet. The bilayer, P_small_, and P_large_ were simulated 2 µs, 5.6 µs, and 5.5 µs, respectively using the Anton 2 supercomputer [58] and the CHARMM all-atom lipid force field [64]. Further build (particularly regarding the fusion pore construction) and simulation details can be found in the Supplemental Material.

### B. Analysis methods

#### 1. The leaflet neutral surface

A key parameter of the analyses is the lipidic *neutral surface* (*δ*), which is the height in a leaflet where curvature and lateral area elasticity are independent. Therefore at *δ*, the area of a leaflet is constant, irrespective of curvature. Curvature-coupled-redistribution (CCR) analysis (see References [41, 43] and the Supplemental Material) indicates that the asymmetric planar bilayer’s *δ* (*δ*_0_, where naught indicates a value for a flat bilayer) to be 13.0 Å (i.e., 14.2 Å for the outer leaflet, 11.7 Å for the inner leaflet). The CCR method works by the fact that the average curvature of an undulating leaflet is only zero when sampled at *δ*_0_. If a surface is chosen above *δ*_0_ (i.e., too close to the lipid headgroups), the surface will appear to have net negative curvature as the headgroup atoms concentrate around a negative (concave) surface. The opposite is true for surfaces defined too close to the midplane. This corresponds with the variation of the area and curvature away from *δ*_0_, as illustrated by Figure 1. Here we assume that *δ*’s position is defined by a particular atom, regardless of leaflet curvature.

**FIG. 1.**
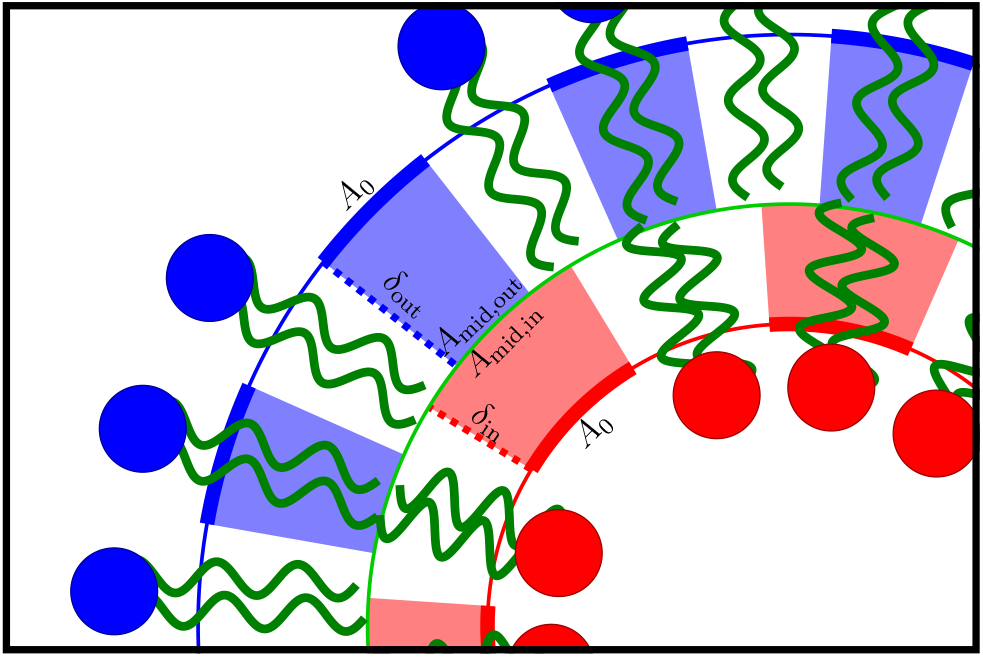
A cartoon showing the variation of thickness in a highly curved bilayer. The *location* of the leaflet *δ*_0_ does not substantially change with curvature, but the hydrocarbon tail *thickness* does change with curvature. Negative curvature (inner leaflet; red) thins the leaflet (changing *δ*_0_ to *δ*_in_, and positive curvature (outer leaflet; blue) thickens the leaflet (changing *δ*_0_ to *δ*_out_). The inner leaflet is thinned more than the outer leaflet is thickened. The length of the heavy red and blue lines are the same, implying area is conserved at *δ*. However, the lengths at the midplane are different (the negatively curved lipid has a larger midplane area (*A*_mid,in_) than the positively curved lipid (*A*_mid,in_)) between the red and blue lipids because of volume conservation [65]. Note that the areas of the red and blue shaded regions are the same.

#### 2. Continuum mesh fits to determine the fusion pore shape

The bilayer and pore shapes were characterized via a fit through their respective midplane surfaces. Using the *δ*_0_ atoms determined by the CCR method, the midplane was fit by matching the continuum surface to the average *δ* collected from the fusion pore simulations. These continuum surfaces contained the contribution from *F*_HC_ as well as a term to favor the surface passing between the leaflet *δ* (Equation S7 of the Supplemental Material).

#### 3. The shape of the pore inner and outer leaflets, given the bilayer midplane surface

In Equation 1, *J*_mid_ and *K*_mid_ are written in terms of the bilayer *midplane*. For strongly curved systems like those considered here, midplane and leaflet curvature differ substantially. The following details how curvature, area, and thickness are determined for *individual lipids* in their respective leaflets, given a description of the bilayer midplane. Consistent with area-preservation at *δ*_0_, we choose to write the lipid area in terms of *δ*_0_’s area, *A*_0_. Figure 1 shows a cartoon of the inner and outer lipid leaflets of a curved bilayer surface. At the bilayer *midplane*, the leaflet areas (*A*_mid,in_ and *A*_mid,out_) are [28]:

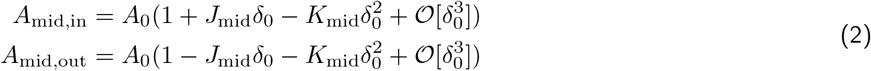

relative to *A*_0_.

The values of *J*_mid_ and *K*_mid_ at the neck for each fusion pore are listed in Table 1. The lightly shaded areas of Figure 1 represent lipid volumes, which are nearly constant with curvature [65]. For the red leaflet, the midplane area (*A*_mid,in_) is larger than *A*_0_. So, to maintain constant volume in the red leaflet, the thickness between the midplane and *δ* decreases.

**TABLE 1.** Basic geometry of the pore neck compared to the bulk.

| Property | Smaller pore | Larger pore |
| --- | --- | --- |
| Neck radius [$\text{\AA}$] | 15.9 | 34.4 |
| Neck $J_{\text{mid}}$ [$\text{\AA}^{-1}$] | -0.01 | 0.013 |
| Neck $K_{\text{mid}}$ [$\text{\AA}^{-2}$] | -0.0017 | -0.0007 |
| Bulk thickness [$\text{\AA}$] | 26.7 | 26.3 |
| Neck thickness [$\text{\AA}$] | 16.2 | 23.0 |
| Neck thinning [%] | 39% | 13% |

The corresponding relations for leaflet interior thickness of the inner and outer leaflets are:

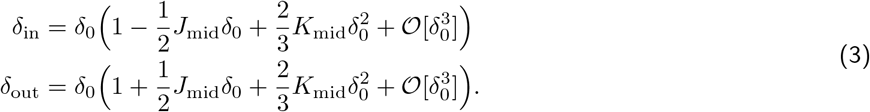

The *bilayer* thickness *δ*_b_ between the neutral surfaces is the sum of the two:

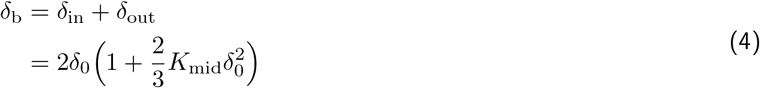

Also see Figure S2 and Supplemental Material for more information on the *K*-induced bilayer thinning. As discussed below, while Equation 4 accurately estimates the bilayer thinning within the large pore’s interior, it underestimates thinning for the smaller pore.

In Figure 1, the radius of the red (*J*_in_ *<* 0) inner leaflet is smaller than the radius of the (*J*_out_ *>* 0) outer leaflet. Leaflet curvatures, *J*_in_ and *J*_out_, where in this model individual lipid curvature energetics area determined, are [8, 66]:

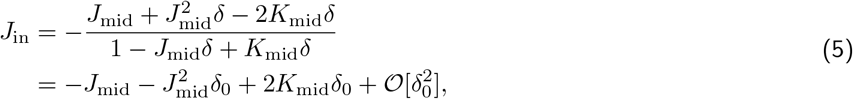

and

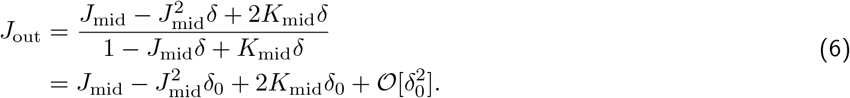

According to Equations 5 and 6, leaflet *J* (*J*_in_ or *J*_out_) changes from *J*_mid_ as a function of *K*_mid_. The change in *leaflet* curvature due to *K*_mid_ can be accounted for in two ways: treating the effect in terms of leaflet curvature or as a change in *κ*_G_. Consider leaflet curvature models of Equation 1 in which *κ*_G_, assumed to be a bilayer rather than leaflet property, is treated separately. Shifting from the midplane to the outer leaflet yields (the bar indicates a *per area* quantity; analysis for 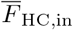 is equivalent):

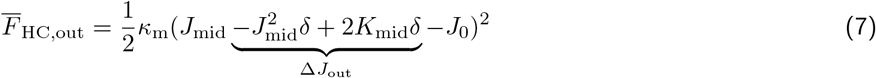

where 2*K*_mid_*δ* is the *K*-based contribution to the shift (Δ*J*_out_). Moving *K*_mid_ out of the squared quantity and dropping higher order curvature terms yields:

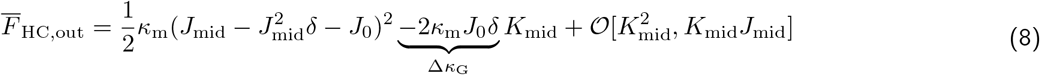

In this form, Δ*κ*_G_ emerges as a leaflet (*κ*_m_ and *J*_0_) contribution to the bilayer treatment of Gaussian curvature. Later in this manuscript, we demonstrate a Δ*κ*_G_ also has a bilayer thickness component that is important for determining fusion pore lipid distributions. The formulation in Equation 8 is important where isolating the effects of *K*_mid_ is important, e.g., in topological changes [8].

## III. LIPID PROPERTIES

To model a lipid’s redistribution on the pore, we employ a local model of elasticity in the vicinity of the lipid. The elasticity energy density 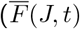; where the bar indicates a density) is:

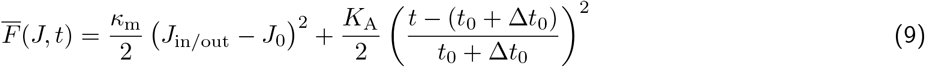

where *κ*_m_ is the leaflet bending modulus 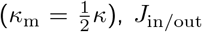 is the total curvature of the inner or outer leaflet, *t* is the measured leaflet thickness, *t*_0_ is the unperturbed, bulk leaflet thickness, Δ*t*_0_ is the difference in thickness preference of an individual lipid from the bulk, and *K*_A_ is the area compressibility modulus (detailed in Reference [67] and the Supplemental Material). Values of *J*_0_ and Δ*t*_0_ are extracted using curvature-coupled relaxation (CCR) and thickness-coupled relaxation (TCR) techniques, respectively, and determined by how lipids redistribute on a planar bilayer’s dynamically fluctuating modes [43]. Cartoons describing the two modes of CCR (left) and TCR (right), as well as the models for chol they imply, are shown in Figure 2 (also see the Supplemental Material for detailed technical information).

**FIG. 2.**
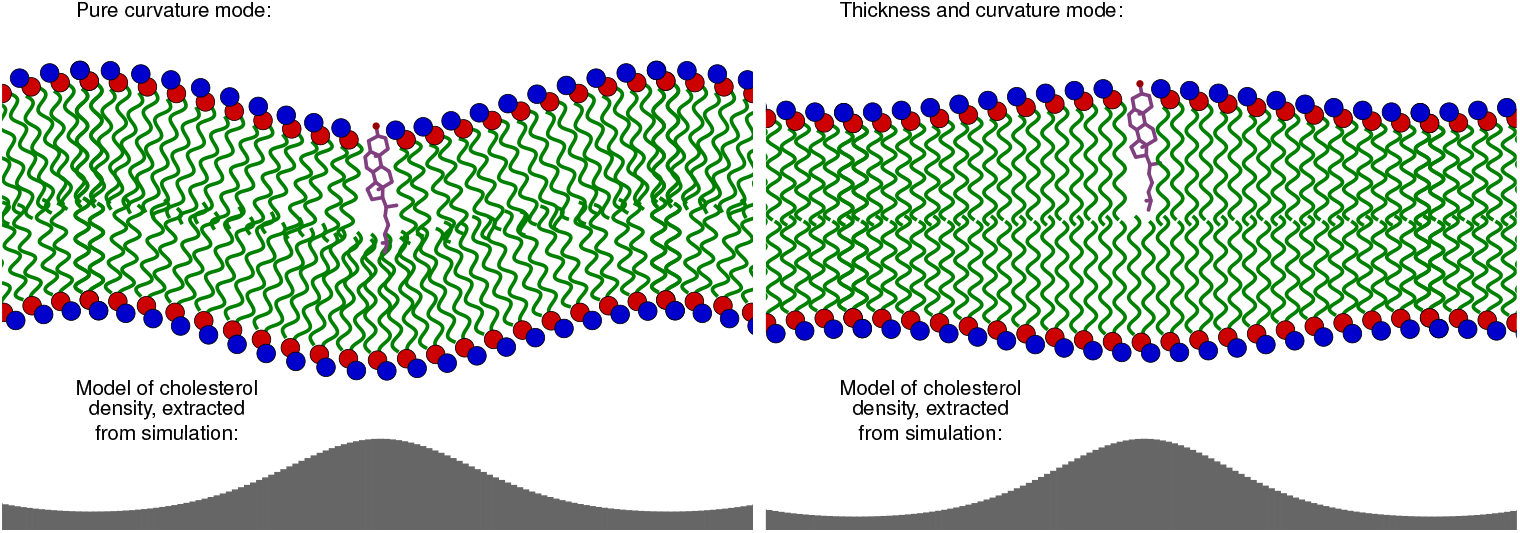
An illustration of the expected distribution of chol according to its curvature (left) and thickness preference (right).

**FIG. 3.**
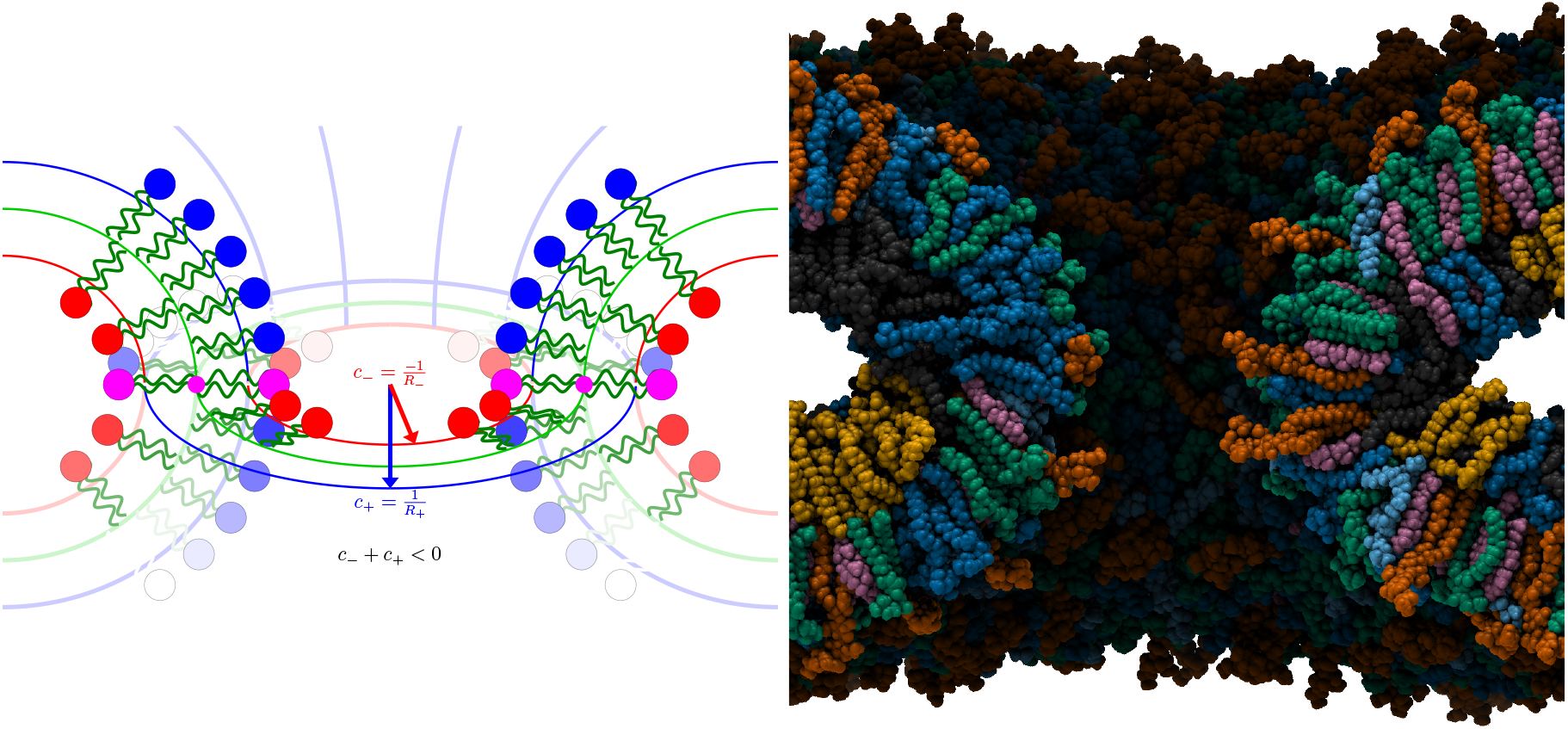
Left: A diagram of curvature on an ideal (*J*_mid_ = 0) fusion pore. At the midplane (small purple dot), the total curvature is zero. At the *leaflet*, total curvature is negative. Right: A molecular image of a snapshot of the large pore (*J*_mid_ = 1.7*×*10^−2^ Å^−1^, with the sign of curvature determined by the outer leaflet).

## IV. GAUSSIAN CURVATURE MODULUS

The Gaussian curvature modulus (*κ*_G_) strongly impacts “topological stability” – the relative free energy of vesicles, sheets, tubes, and strange objects like the cubic phase [8, 9, 30] and cubosomes [68]. This is because, unlike the curvature *J*, the integrated Gaussian curvature on a surface, *∫* _*S*_ *K*, is a constant that only depends on topology via the Gauss-Bonnet theorem. Although a full mechanistic understanding of *κ*_G_ is unknown, differences between lipids are primarily understood through its dependence on *J*_0_ as in Equation 8. In this work, we are able to extend the treatment of the lipid-specific dependence of *κ*_G_ by incorporating the thickness elasticity of the bilayer interior.

According to Equation 4, a bilayer with saddle curvature (*K <* 0) experiences *net thinning* within the thickness of the bilayer interior, *t*_0_ = 2*δ*_0_. The strain is equal to:

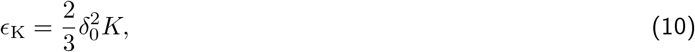

where *ϵ*_K_ is the strain away from the total bilayer thickness. The *individual* lipid strain *ϵ*_l_ is

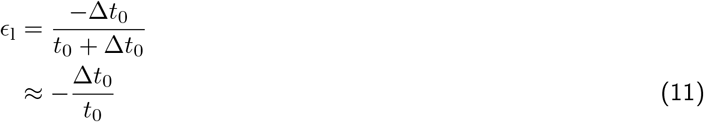

where Δ*t*_0_ is the difference in bilayer thickness preference of the lipid, compared to *t*_0_. The numerator is the deviation from the preferred thickness (*t*_0_ + Δ*t*_0_). To first order, strains are additive:

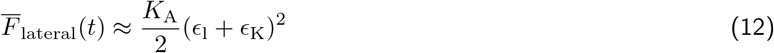

Expanding the local elastic thickness energy in powers of *K* yields:

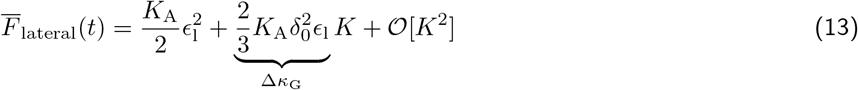

The first energy term in Equation 13 is from the thickness strain (dependent on the compressibility modulus, *K*_A_), and the second term is from the Gaussian curvature strain (dependent on this component of the Gaussian curvature modulus, Δ*κ*_G_). The predicted differences in bilayer *κ*_G_ for each lipid, derived from bilayer thickness elasticity, are listed in Table S6. These values are derived from *ϵE*_l_ in Equation 11 multiplied by 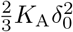 :

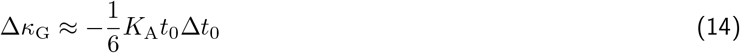

with *t*_0_ = 2*δ*_0_. Note that Δ*κ*_G_ is inherently a *bilayer* property, as it is determined by the interior thickness of a bilayer. Contrast this with the influence of lipid spontaneous curvature on *κ*_G_ (Equation 8) which is solely an effect of leaflet spontaneous curvature.

## V. LIPID REDISTRIBUTION FROM ALL-ATOM SIMULATION AND LITERATURE ELASTICITY THEORY

In the simplest version of the HC model, lipids have a single spontaneous curvature *J*_0_ that does not depend on the context of the surrounding leaflet (e.g., the composition). Further assuming the effect of a lipid *i* is localized [43], its contribution to the bending free energy (*F*_i_) is:

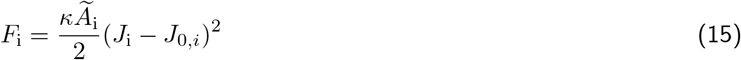

where *κ* is the bending modulus, *Ã*_i_ is lipid *i*’s lateral area, *J*_*i*_ is the curvature local to a lipid *i*, and *J*_0,*i*_ is lipid *i*’s intrinsic curvature. The likelihood *p*_i,r_ of observing the lipid in a particular region *r* with curvature *J*_r_ is proportional to its Boltzmann factor:

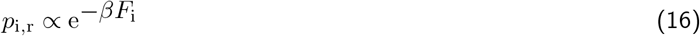

but naturally will be influenced by similar energetics of the other lipids. Therefore, lipid populations in each region are predictable.

Here, we use a Monte Carlo (MC) model to predict equilibrium lipid distributions on the fusion pores. Initially, the MC simulations are informed by *J*_0_, *Ã, κ*, and *K*_A_ that are determined either by previously published simulation or by analogy. The MC allows us to make specific hypotheses on where a particular lipid will reside can be tested based on current literature expectations and assumptions. Imperative to this paper, the MC provides further evidence that Δ*κ*_G_ strongly determines lipid distributions in small fusion pores (see below). The MC details, including Table S7 showing mechanical parameters, can be found in the Supplemental Material.

## VI. RESULTS AND DISCUSSION

### A. Lipid redistribution timescale in the fusion pore neck

Lipids are expected to diffuse across the neck with timescale *q*^*−*2^*D*^*−*1^, where *D* is the diffusion constant, *q* = 2*π/λ* and *λ* is the lengthscale (*∼*8 nm) of the region. With *D ≈* 8 µm^2^/s, the composition is expected to relax in *∼*200 ns. Statistically, the variance of chol’s mol fraction in the region is *σ*^2^ = *p*(1 *− p*)*/n*, where *p* is the mol fraction of chol and *n* is the number of lipids in the region. For a chol mol fraction of 0.30 (i.e., *ϕ*_chol_ = 0.30) and *∼*100 lipids in the neck of P_small_, *σ* = 0.046. In P_large_, *n* = 150 and *σ* = 0.037. Figure 4 shows the timeseries of *ϕ*_chol_ in the small pore over the course of the simulation, with the autocorrelation function inset. Although the autocorrelation is not sufficiently converged to extract a diffusion constant, it is useful for a rough indication of the timescale. Both the magnitude of the fluctuations (which are damped by sampling over a 24 ns window) and timescale of relaxation are both consistent with our estimates.

**FIG. 4.**
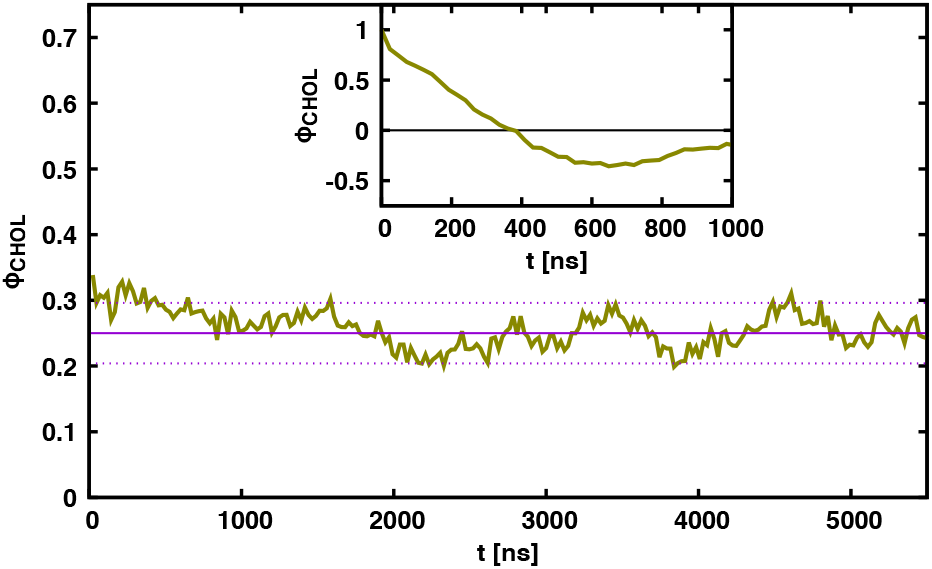
Timescale of the chol fraction (*ϕ*_chol_) in the neck of the small pore. The mean over the last 4.5 microseconds is shown with a solid purple line. Dotted lines indicate one standard deviation expected from statistical fluctuations. Instantaneous fluctuations were reduced by computing the composition over 24 ns windows. The autocorrelation function (shown in the inset) is consistent with a _*∼*_200 ns timescale for compositional relaxation.

### B. Curvature elasticity partially describes the observed lipid redistribution

Following *∼*5.5 µs simulations, the lipid distributions for P_small_ and P_large_ were calculated (solid bars in Figure 5). For P_small_, the outer leaflet shows strong enrichment of PLPC and PAPC with depletion of GM3, PSM, and chol. Most of the redistribution results can be described by *J*_0_: unsaturated PC lipids have a negative curvature preference [69] and sphingolipids have a positive curvature preference [70]. In the inner leaflet, SAPE is enriched, SAPS is near the bulk value, and PLPC and chol are depleted. The MD results suggest that SAPE simply out-competes PLPC and SAPS because of a strongly negative *J*_0_. However, with its strong negative curvature, chol should be enriched in the neck to a similar degree as SAPE.

**FIG. 5.**
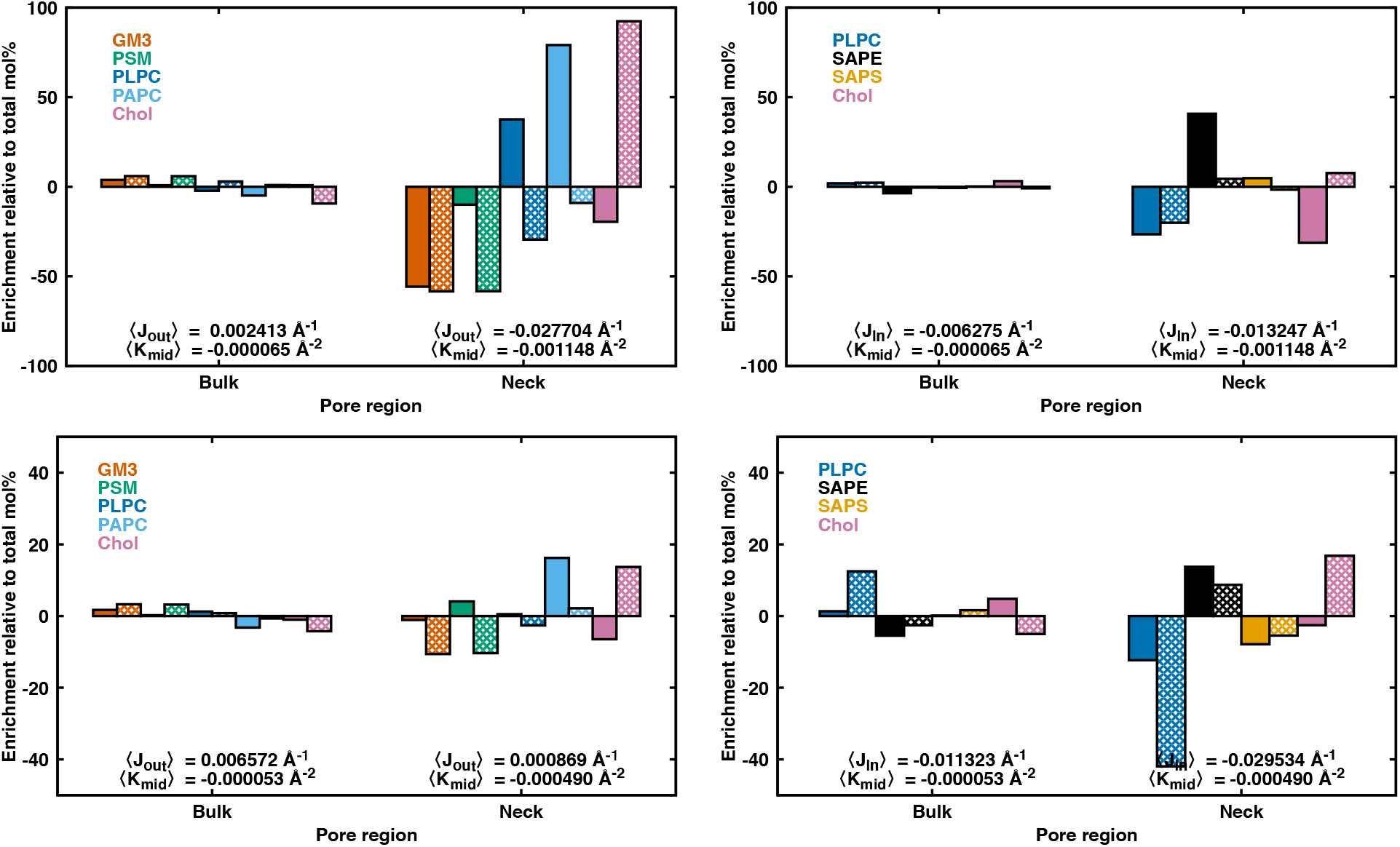
Enrichment per region relative to the total mol% of a lipid species using only *J*_0_, *κ*, and area contributions (enrichment = 100 *×* (*ϕ*_species,observed_ *− ψ*_species,total_)*/ψ*_species,total_). Data for the small pore’s outer (left) and inner (right) leaflets on the top row, respectively. Data for the large pore’s outer (left) and inner (right) leaflets on the bottom row, respectively. See Table S1 for *ϕ*_species,total_ values. Direct observations from MD are solid bars and MC predictions are hatched. The MC predictions do not include a Δ*κ*_G_ contribution.

To gain further energetic insight into these results, we utilized a MC model using only *J*_0_, *κ*, and area contributions to the free energy (i.e., no Δ*κ*_G_; see Equation S36 of the Supplemental Material). The results are shown as hashed bars in Figure 5. MC predicts depletion of GM3 and PSM based on their positive *J*_0_ and a depletion of PLPC based on competition. Importantly, the MC model predicts strong chol enrichment given its very strong negative *J*_0_. In the inner leaflet, SAPE and chol are expected to be enriched in the pore. However, chol is depleted in the MD. Similar results are shown for P_large_. Therefore, a simple model containing only information on *J*_0_, *κ*, and area contributions does not adequately capture the complex energetics occurring in the MD. The stark mismatch between the MC and MD suggests that there are more complex curvature energies to be determined.

### C. Gaussian curvature thins the bilayer interior in the pore and rejects chol

Applying Equation 4 to the midplane saddle curvature in Table 1 yields the expected thinning in the bilayer interior (*δ*_b_) at the most extreme curvature of the pore. For the small pore, the expected thinning is 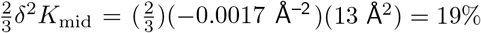, while for the large pore it is 8%. Simulations indicate that thinning for the small and larger is even more dramatic, at 39% and 13%, respectively (Figure 6). Two possibilities may explain the discrepancy. First, chol exclusion from the neck would thin the bilayer, i.e., composition alone accounts for the thinning. To test this, planar bilayers were built with asymmetric compositions equivalent to those of the pores. Based on thickness analysis of these planar bilayers, the necks of P_small_ and P_large_ should have *δ*_b_ similar to the bulk, far from the pore’s neck. Therefore, the observed thinning in the neck cannot be explained simply by local composition. See the Supplemental Material for more information on these simulations and analyses. Second, thinning is sensitive to the precise location of *δ*, going as the square of *δ*. Higher order effects and deformations inconsistent with local volume conservation may shift the precise location of *δ*. Figure S2 in the Supplemental Material shows the variation of thinning with pore radius, as a function of *δ* – large variations in thinning occur with small shifts of *δ* or the pore radius.

**FIG. 6.**
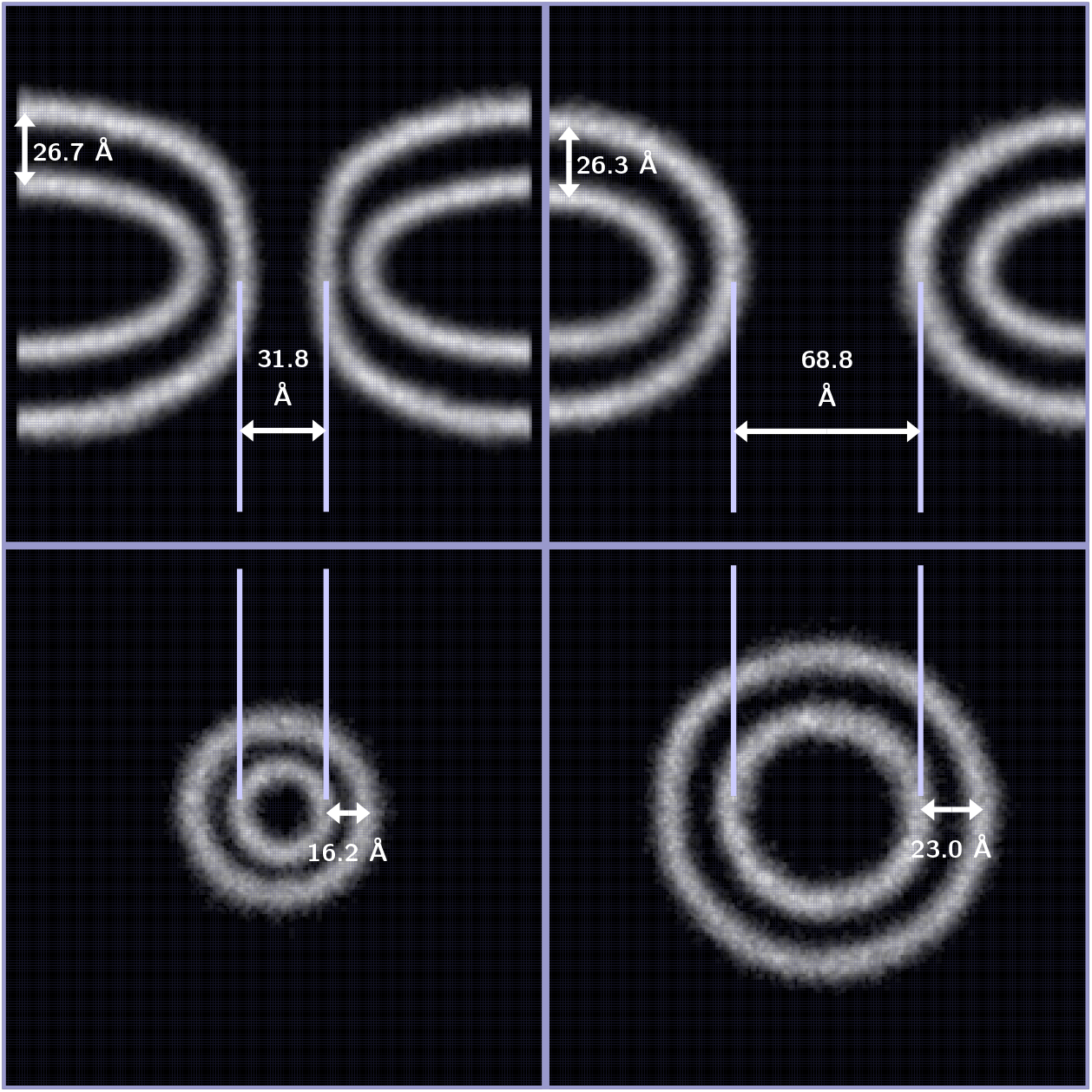
The density of *δ* atoms for a 5-Å-wide slice. Left column: the smaller pore. Right column: the larger pore. The top row is a 5-Å-wide slice in *xz* slice through *y* = 0. The bottom row is a 5-Å-wide slice in *xy* through the pore at *z* = 0. Each box is 200 Å by 200 Å.

**FIG. 7.**
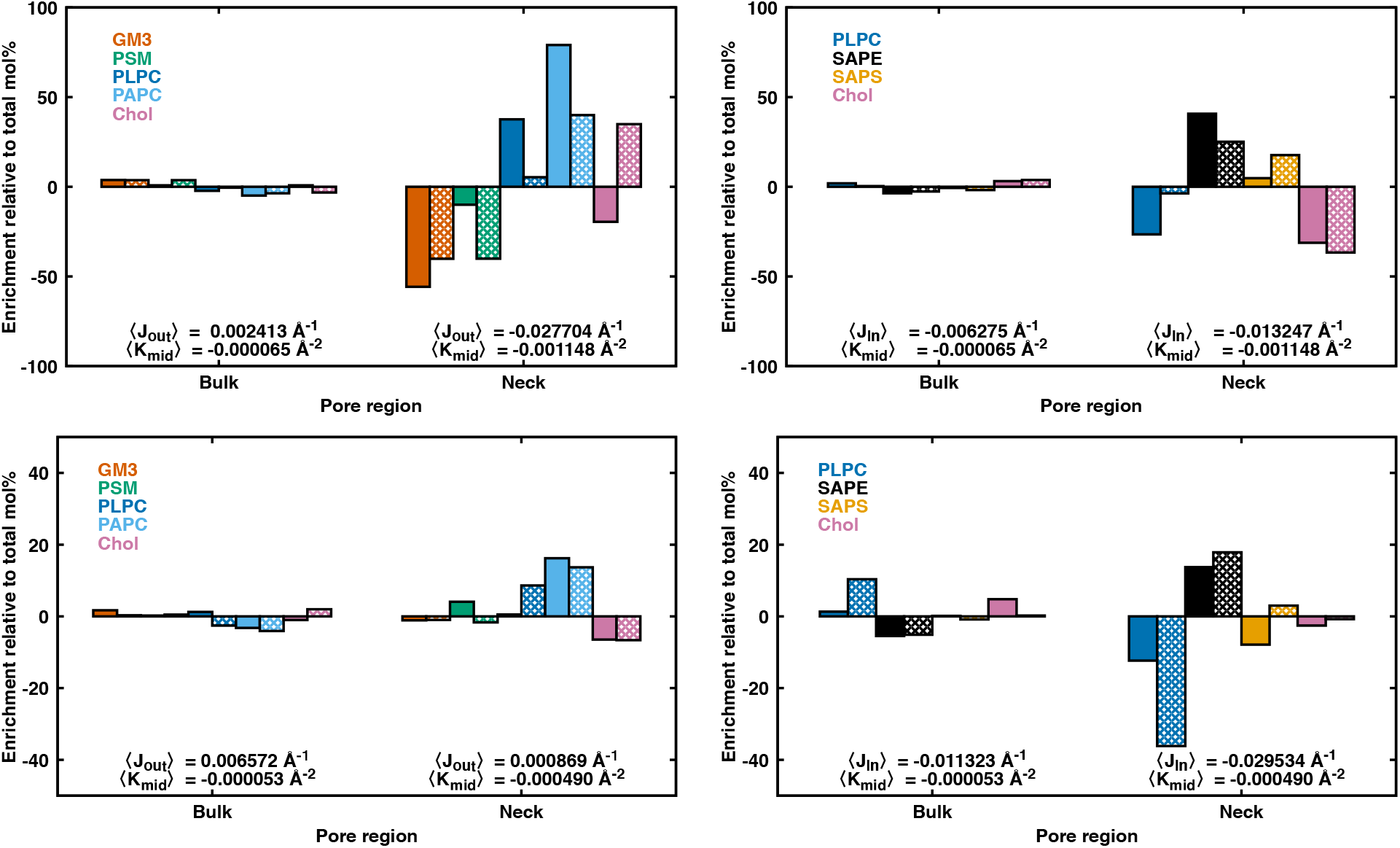
Enrichment per region relative to the total mol% of a lipid species including Δ*κ*_G,chol_ (enrichment = 100 *×* (*ϕ*_species,observed_ *− ψ*_species,total_)*/ψ*_species,total_). Data for the small pore’s outer (left) and inner (right) leaflets on the top row, respectively. Data for the large pore’s outer (left) and inner (right) leaflets on the bottom row, respectively. See Table S1 for *ϕ*_species,total_ values. Direct observations from MD are solid bars and MC predictions are hatched.

Thinning in the bilayer interior does not necessarily indicate total (e.g., headgroup to headgroup) thinning. Indeed the same considerations leading to thinning in the interior lead to thickening in the exterior (head group region). This is the basis for the long-established theory for why lipids with negative *J*_0_ favor cubic phases [8]. Yet the theory is inconsistent with chol. Motivated by the fact that chol resides mostly within the bilayer interior, we hypothesize that chol is sensitive to interior thinning to the same degree it is sensitive to total bilayer thinning.

### D. Adding Δ*κ*_G_ into the Helfrich-Canham (literature) model improves the description of lipid redistribution

Given bilayer thinning from saddle Gaussian curvature is an important geometric constraint on these fusion pores, we introduced Δ*κ*_G,chol_ into the MC model (Equations 1 and S36–S37). Δ*κ*_G_ was calculated using the TCR method on the planar bilayer simulation (see Supplemental Material Figure S5 and Table S6). Similar to the CCR method that obtains *q*-based *J* preferences for a given lipid species, TCR obtains *q*-based *thickness* preferences. Using Equation 14, we can calculate the bilayer thinning contribution to the Gaussian bending modulus (Δ*κ*_G_). Most striking is that Δ*κ*_G,chol_ = –10.4 *±* 3.5 kcal/mol in the fluid inner leaflet (this value is reduced to –4.7*±* 2.2 kcal/mol in the outer leaflet). A lipid with negative Δ*κ*_G_ will be repelled from the pore where there is strong negative (saddle) *K* (Equation 1). We test the hypothesis that only chol is strongly influenced by Δ*κ*_G_ because it is the only lipid that nearly-fully resides in the bilayer hydrophobic interior, and therefore, could be especially prone to bilayer thinning. We use the inner leaflet’s Δ*κ*_G,chol_, assuming that the lipids are in a disordered state. By simply introducing Δ*κ*_G,chol_ into the MC model, all enrichments/depletions in both leaflets of the P_small_ and P_large_ are much better fit relative to the MC model without Δ*κ*_G,chol_ (7). Still, the largest discrepancies occur in P_small_’s outer leaflet, which has the strongest curvature considered in these simulations.Using Δ*κ*_G,chol_ = –13.9 kcal/mol (i.e., one standard error away from the mean) further improves the MC fit to P_small_’s outer leaflet MD data (see Figure S7). We conclude that Δ*κ*_G,chol_ is an imperative energetic term for determining the lipid distributions of small fusion pores.

### E. Evolution of fusion pore shape

Given the complexity of the fusion pore shape, we explore two sources of stress: i) hydrostatic/osmotic stress caused by having two distinct water compartments, and ii) differential stress caused by having too many/few lipids per leaflet. We assess how these stresses affect the simulation results.

#### 1. Relieving hydrostatic stress

Because of the fusion pore’s shape and periodic boundary conditions, there is a trapped water compartment in which the number of water cannot change (also see Reference [57]). Water is incompressible relative to the membrane, so the fixed amount of water determines the membrane shape. To test the ramifications of our initial conditions, we added carbon nanotubes (CNTs) into the membrane bulk and simulated a further *∼*1.5 µs using Amber [71, 72] and the CHARMM all-atom force field (see the Supplemental Material for detailed simulation methodology). In P_small_, water exits the compartment, whereas in P_large_, water enters the compartment. The water flow in the systems enables the radius of P_small_ to expand and for the P_large_ to contract (see Figures S8 and S9). This indicates that our fusion pore simulations without CNTs were under hydrostatic stress, but *∼*1.5 µs with CNTs is insufficient to fully assess how the hydrostatic force affected the fusion pore shape.

#### 2. Leaflet differential stress

Differential stress in a bilayer occurs, for example, when there are too many or too few lipids in a leaflet (i.e., an area imbalance). This stress can be relieved by bending and flipping chol. As discussed in the previous section, without CNTs, the pore geometry is fixed. Therefore, we quantify chol flips to assess the differential stress. Chol flips were identified by comparing chol orientation normal to the continuum fit surface. Identified flips lasting *<* 40 ns were filtered to avoid including assignment errors. We found that in the pores, chol flips between leaflets at a sub-millisecond rate (*∼*0.1–0.2 ms^*−*1^), a rate which was roughly similar across all four simulations (P_large_ = 0.14 ms^*−*1^; P_small_ = 0.15 ms^*−*1^; P_large, CNT_ = 0.19 ms^*−*1^; P_small, CNT_ = 0.13 ms^*−*1^). Flips tended to occur near the pore neck where the bilayer is thinner, however, P_small_, with its substantially thinner neck, had a similar number of flipping events as P_large_. The rates are similar to previous observations of flip-flop in heterogeneous bilayers, in which the ordered phase timescale (0.1 ms^*−*1^) was slower than that of a completely fluid phase (1.0 ms^*−*1^) [73]. A rough comparison suggests that the flipping events in the relatively small part of the neck, where a minority of chol are located, are occurring at fluid-phase timescales, while transitions in the bulk are much slower, comparable to the ordered-phase timescale. However, targeted methodology is necessary to statistically differentiate the rate of flip with pore size, composition, and localization. See the Supplemental Material for a schematic showing locations of the chol flips on the pores (Figure S8) and a further discussion on differential stress.

### F. Consequences for endo-/exo-cytosis

All-atom MD and a literature-based MC simulation indicate chol favors thicker membranes and is therefore excluded from thin fusion pores. Therefore, we ask how a simple model predicts the energetics of pore closure as a function of chol exclusion. The model used herein is related to that of Chen, Higgs, and MacKintosh, who determined how a minority membrane component can initiate pore collapse leading to fission [59]. Our model focuses on the entropic cost of chol depletion in the fusion pore and the neck’s thickness strain (Equation 4). First, entropically, it is easier to deplete a small region of all chol than a large region, therefore, the cost of chol depletion increases with fusion pore size. Second, the *total* thickness strain is constant in the absence of redistribution because *K* is a constant that only varies with topology (i.e., area strain is proportional to ∫_*S*_ *K*). This effect is borne out in the following model, which is treated in terms of chol but is applicable to arbitrary lipids.

Chol’s free energy contribution to the pore (*F*_pore_) is modeled with two terms, *F*_pore_ = *F*_entropy_ + *F*_HC_. First is the entropy of the mol fraction (*ϕ*) of chol in the vicinity of the fusion pore (*F*_entropy_):

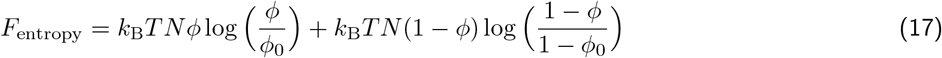

where *F*_entropy_ arises simply from counting the ways of arranging chol and the other lipids on a surface accommodating *N* total lipids (i.e., *Nφ* is the number of chol in the region). The free energy is minimized when *ϕ* equals the total leaflet mol fraction *ϕ*_0_. Expanding *F*_entropy_ around *ϕ*_0_ (*ϕ* = *ϕ*_0_ + Δ*ϕ*) yields:

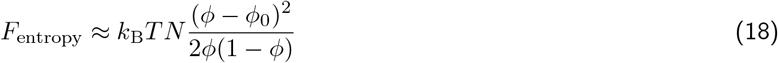

That is, there is an effective force constant 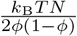 restricting fluctuations around *ϕ*_0_. As *N* grows (proportional to the size of the pore), the force constant becomes more restrictive.

Second is chol’s HC energy in the neck relative to the bulk (*F*_HC_; considering the average *J* and *K* of the region, ⟨ *J* ⟩ and ⟨ *K*⟩, respectively):

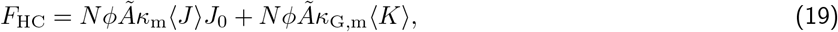

The *F*_HC_ term is similar to the Monte Carlo method for determining lipid enrichment, described in detail in the Equation S36. The shape, and thus ⟨ *J* ⟩, of the ideal fusion pore will depend on the lipids, their *J*_0_, as well as the protein system (e.g. clathrin and dynamin) that is forcing the constriction. For simplicity here we only model the effect of ⟨ *K* ⟩ on the free energy (*F*_chol,saddle_), equivalent to setting ⟨ *J* ⟩ = 0:

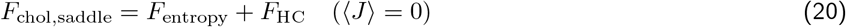

With *N* ⟨ *K* ⟩ = ∫ _*S*_ *K* = *−*4*π* (the topological invariant for the saddle simulated here, see Figure 2 of Reference [8]):

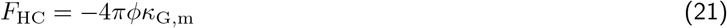

Minimizing *F*_chol,saddle_ with respect to *ϕ* to obtain *ϕ*^*∗*^ (the optimal *ϕ*), yields:

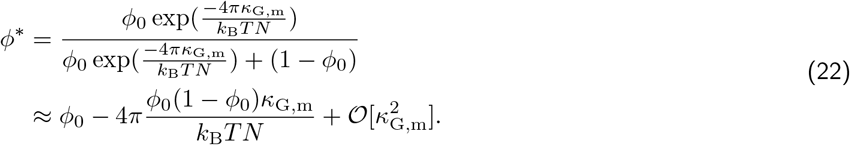

That is, deviations from *ϕ*_0_ are damped as the pore size grows, and conversely, a shrinking pore (i.e., small *N*) allows larger deviations from *ϕ*_0_.

Inserting *ϕ*^*∗*^ from Equation 22 into *F*_HC_ + *F*_entropy_ yields:

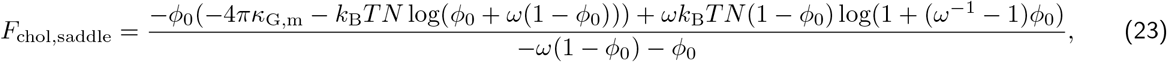

where for clarity we have defined 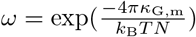. Its asymptote for large *N* is:

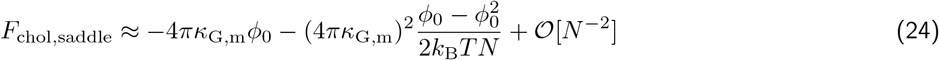

and its asymptote for small *N* is:

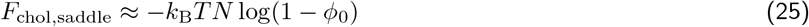

Note that Equation 25 requires *κ*_G,m_ to deviate sufficiently from zero such that redistribution of the target lipid into or out of the pore is nearly complete for small pores.

To discuss the model in terms of fusion pore size instead of numbers of lipids, we assume a rough scaling of the fusion pore’s radius (*r*) with *N* :

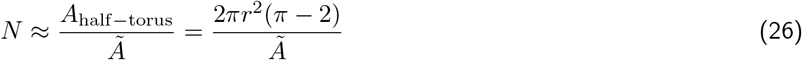

where *A*_half*−*torus_ is the inner-half area of a torus with saddle curvature *K* = *− r*^*−*2^ at the inner rim. Variations on Equation 26 from a half-torus will change the details of the free-energetic force to collapse the pore, but will not change the energetic scale, which asymptotes to *−*4*πκ*_G,m_*ϕ*_0_ regardless of how the area scales with *N*.

Figure 8 shows the variation of the free energy due to chol redistribution as a function of approximate fusion pore size. As *r* decreases, the entropic penalty (Equation 18) for depleting chol from the fusion pore decreases, while the elastic benefit for depletion per unit *ϕ* (Equation 21) is constant. The asymptote in Equation 24 (Figure 8, blue dashed line) describes the variation in free energy well for nearly the entire region of interest before pore collapse. Once chol is nearly completely depleted from the pore, the free energy continues to drop only due to lesser entropic penalty (red dashed line, Equation 25). At this point the fusion pore radius is so small that an elastic analysis is likely inappropriate.

**FIG. 8.**
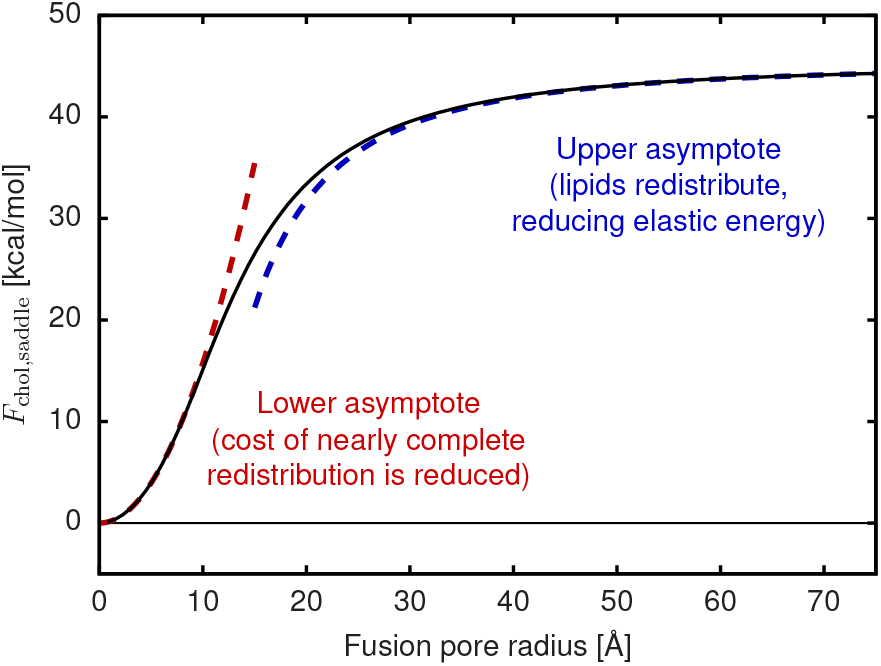
A plot of the modeled effect of chol redistribution on the fusion pore energy (Equation 23). The large *N* (Equation 24) and small *N* (Equation 25) asymptotes are shown.

According to this simple model, a chol-enriched bilayer (or any multi-component system with varied *κ*_G_, as originally predicted in Ref. [59]) will favor the transition from a flat clathrin plaque to a bud, with the bilayer appearing to be softer than its *κ* might imply. This effect belongs to a general class of softening mechanisms for multicomponent systems [41, 74– 77]. We believe this is the mechanical origin of the requirement for sterols in clathrin-mediated endocytosis observed in Ref. [16].

## VII. CONCLUSIONS

All-atom molecular dynamics simulations of two fusion pores indicate substantial lipid redistribution between the pore and bulk, depending on pore size. Continuum mesh surfaces were fit to the pores to determine the total (*J*) and Gaussian (*K*) curvature in the bulk and neck region of the pores. Comparing the lipid distributions from simulation to the literature Helfrich/Canham model, we found that lipid redistribution cannot be adequately described using the spontaneous curvature (*J*_0_) and bending modulus (*κ*_m_) alone. We explain the anomalous depletion of chol from the pore neck on the basis of its Gaussian curvature modulus, *κ*_G_. Strong saddle curvature thins the *bilayer* interior (i.e., not just a single leaflet), which disfavors chol. Dynamic redistribution of chol on planar simulations of similar composition yielded estimates for the thickness-driven change in Gaussian curvature modulus (Δ*κ*_G_) for each lipid in the bilayer simulation. Importantly, chol in the fluid inner leaflet has a strong negative Δ*κ*_G_ (–10.4 *±* 3.5 kcal/mol), indicating a lipid that is repelled by saddle curvature. Using this information, we informed the literature model to include the *K* and Δ*κ*_G_ observed in the MD simulations. This addition improved the theoretical model’s fit to the simulation data, indicating that *K* and Δ*κ*_G_ determine lipid distributions during fusion pore opening/closure. In the smaller pore, ganglioside GM3, palmitoylsphingomyelin (PSM), and chol were all excluded from the fusion pore’s neck – suggesting a structural connection to L_o_ domain formation that would likely further alter lipid material parameters. Theoretical consideration of a two-component system (e.g., chol and the surrounding lipid matrix) demonstrated that it is energetically easier to deplete chol from smaller structures, such that chol is depleted to a much greater extent in smaller fusion pores. This is observed in the all-atom molecular dynamics simulations. We hypothesize that *K*-driven chol redistribution in fusion and fission pores favors pore closure. Therefore, increased chol levels should favor the late stages of endocytosis and budding (which require closure), but disfavor viral fusion, which requires fusion pore expansion.

## VIII. ACKNOWLEDGMENTS

This work was supported by the Intramural Research Program of the *Eunice Kennedy Shriver* National Institute of Child Health and Human Development (NICHD) at the National Institutes of Health. A.H.B. was supported by a Postdoctoral Research Associate (PRAT) fellowship from the National Institute of General Medical Sciences (NIGMS), award number 1Fi2GM137844-01. Anton 2 computer time was provided by the Pittsburgh Supercomputing Center (PSC) through Grant R01GM116961 from the National Institutes of Health. The Anton 2 machine at PSC was generously made available by D.E. Shaw Research. Initial setup and analyses were performed on computational resources provided by the Intramural Research Program of the NICHD. We thank Hai Lin for helpful discussions on equilibrating osmotic forces on the pore. Molecular rendering was performed with Tachyon software written by John E. Stone.

## IX. CONTRIBUTIONS

A.H.B. and A.J.S. developed the approach, designed the system setup, and performed the simulations. All authors performed the analysis and wrote the manuscript.

