## Supplemental Material for "Dynamic cholesterol redistribution favors membrane fusion pore constriction"

### I. SYSTEM CONSTRUCTION AND SIMULATION

#### A. Continuum modeling for fusion pore construction

A key advance of this manuscript is the use of continuum meshes to both build and analyze the fusion pores. The basis of the mathematical description of the surface by a set of vertices connected to form a triangular mesh. The tangent-plane continuous surface is defined via the subdivision-limit algorithm [1], developed for use for membranes by Feng and Klug [2].

Formally, a limit surface is determined by iterating a subdivision operation on the triangles of a mesh. The triangles of a mesh at iteration $i$ are each divided into four smaller triangles by dividing the edges of the triangle in half, with the mid-points forming an interior triangle. The vertices of the next iteration are determined by the rule:

$$\boldsymbol{r}_{a,i} = \frac{5}{8}\boldsymbol{r}_{a,i-1} + \sum_{b\in a}^{N} \frac{3}{8N}\boldsymbol{r}_{b,i-1} \tag{S1}$$

$$\boldsymbol{r}_{\text{new},i} = \frac{3}{8}(\boldsymbol{r}_{\mathit{1},i-1} + \boldsymbol{r}_{\mathit{2},i-1}) + \frac{1}{8}(\boldsymbol{r}_{\mathit{3},i-1} + \boldsymbol{r}_{\mathit{4},i-1}) \tag{S2}$$

where $N$ is the valence of a vertex, $b \in a$ refers to the set of vertices with edges to $a$, indices $\mathit{1}, \mathit{2}$ refer to the vertices of the subdivided edge, and $\mathit{3}, \mathit{4}$ refer to the two vertices of the triangles $\mathit{123}$ and $\mathit{124}$ that share the edge $\mathit{12}$. Infinite application of the rule yields a set of points that constitute a $C^1$ continuous surface [3].

The mathematical form of the surface is a cubic spline [1] that is a function of the starting vertices, termed the *control points*. The Helfrich-Canham (HC) energy can then be minimized by optimizing the positions of the control points. The resulting surface can be used to evaluate the curvature everywhere except at a discrete set of points at irregular vertices (vertices with more or less than six edges). Practically this is irrelevant – the energy is typically integrated over a finite set of points on the surface that does not include the problematic point. For regular vertices we use fifteen points distributed equally across the triangle. For irregular vertices, we use sixteen-point Gaussian integration optimized for the form of the HC energy on an irregular face.

#### B. Main all-atom bilayer and fusion pore construction/equilibration details

Leaflet concentrations for the bilayer, $P_{small}$, and $P_{large}$ (Tables S1–S3)

| Leaflet | chol | GM3 | PSM | PLPC | PAPC | SAPE | SAPS |
|---|---|---|---|---|---|---|---|
| Outer | 0.30 | 0.10 | 0.28 | 0.25 | 0.07 | — | — |
| Inner | 0.30 | — | — | 0.16 | — | 0.34 | 0.20 |

TABLE S1. Initial bilayer leaflet concentrations. There were 451 lipids in the outer leaflet and 400 lipids in the inner leaflet. Chol flip-flop slightly changes the leaflet compositions.

| Leaflet | chol | GM3 | PSM | PLPC | PAPC | SAPE | SAPS |
|---|---|---|---|---|---|---|---|
| Outer | 0.32 | 0.09 | 0.29 | 0.24 | 0.07 | — | — |
| Inner | 0.30 | — | — | 0.17 | — | 0.31 | 0.22 |

TABLE S2. Initial $P_{small}$ leaflet concentrations. There were 1628 lipids in the outer leaflet and 1353 lipids in the inner leaflet. Chol flip-flop slightly changes the leaflet compositions.

| Leaflet | chol | GM3 | PSM | PLPC | PAPC | SAPE | SAPS |
|---|---|---|---|---|---|---|---|
| Outer | 0.33 | 0.10 | 0.29 | 0.23 | 0.06 | — | — |
| Inner | 0.32 | — | — | 0.17 | — | 0.32 | 0.19 |

TABLE S3. Initial $P_{large}$ leaflet concentrations. There were 1895 lipids in the outer leaflet and 1319 lipids in the inner leaflet. Chol flip-flop slightly changes the leaflet compositions.

##### 1. All-atom bilayer construction and equlibration

Symmetric bilayers of "inner" or "outer" composition were built [4–7] and simulated 50 ns with NAMD [8, 9] and the CHARMM all-atom lipid force field [10] to obtain the equilibrium leaflet area of both compositions. Once the equilibrium area of both symmetric patches was quantified, the inner and outer leaflet compositions were scaled to construct an asymmetric bilayer presumably without latent stress [11, 12]. See **Table S1** for lipid compositions. The asymmetric bilayer was neutralized, and the solution was brought to ~150 mM KCl. After a 20 ns equilibration using NAMD [8, 9], the system was converted to dms format using the `charmmgui2dms.py` code written by Yifei Qi and available on the CHARMM-GUI server [5]. The NAMD simulations used a temperature of 310.15 K set by a Langevin dynamics piston (1 $ps^{-1}$ damping coefficient), a semi-isotropic pressure of 1 atm set by a Nosé-Hoover Langevin piston (50 fs period and a 25 fs decay time) [13], a 2 fs timestep, and covalent bonds involving hydrogen were constrained by SHAKE and SETTLE [14, 15]. Non-bonded interactions were switched off between 10–12 Å, and electrostatics were calculated by smoothed particle mesh Ewald (PME) with a maximum of 1 Å between grid points [16, 17].

##### 2. Fusion pore construction/equilibration details

Fusion pore geometries were built by connecting two planes with a cylinder, and the continuum mesh was minimized with respect to the HC energy. This mesh defined the bilayer midplane. To obtain the distorted, larger pore, the volume inside the inner compartment was constrained to match a target initial condition. The atomistic fusion pores were built by transforming patches of a asymmetric, planar "source" simulation into the curvature and orientation of the "target" fusion pore continuum model. The procedure assumes a standard mechanical leaflet deformation to match the local spacing and orientations of the source and target lipids. The key parameter is the same neutral surface ($\delta$) described in the main text: the plane defined by a distance $\delta$ along the normal from the bilayer midplane where area and curvature deformations appear to be uncoupled. The area-per-lipid is constant at $\delta$ as the leaflet curves. In short, the triangles of the target mesh were broken up into contiguous regions that use the same source-to-target transformation. In this way, much of the curved leaflet has similar packing to the planar leaflet, with the exception of the boundaries between regions.

For each region on the target pore, a source point was randomly chosen in the planar simulation (centered on a lipid) and a corresponding point was chosen in the center of the target region. For the source point, the $x, y, z$ coordinate system of the planar system, with $z$ approximately along the normal was mapped to a rotated and translated Cartesian system at a point on the pore, $\{x', y', z'\}$. At the target point, the *curvilinear* coordinate system was used to determine the position of the target lipid. The planar $z$ direction matches the surface normal on the pore ($z'$). In this way, the planar coordinate system $\{\boldsymbol{x}, \boldsymbol{y}, \boldsymbol{z}\}$ was mapped to a new coordinate system $\{\boldsymbol{x}', \boldsymbol{y}', \boldsymbol{n}\}$, where $\boldsymbol{x}'$ and $\boldsymbol{y}'$ are orthonormal at the target point, and depend on the surface coordinates $u$ and $v$. The Cartesian axes in the pore frame are mapped to the subdivision coordinates $u$ and $v$ that define $\boldsymbol{r} \equiv \boldsymbol{r}(u, v)$:

$$\boldsymbol{x}' = \frac{\partial \boldsymbol{r}}{\partial u} x_u + \frac{\partial \boldsymbol{r}}{\partial v} x_v \quad \text{(S3)}$$

$$\boldsymbol{y}' = \frac{\partial \boldsymbol{r}}{\partial u} y_u + \frac{\partial \boldsymbol{r}}{\partial v} y_v \quad \text{(S4)}$$

where scalar constants $x_u$, $x_v$, $y_u$, and $y_v$ are chosen such that $\boldsymbol{x}'$ and $\boldsymbol{y}'$ are orthonormal. Rotation of $\boldsymbol{x}'$ and $\boldsymbol{y}'$ is arbitrary and chosen randomly; however, the same rotation is chosen for each lipid in the patch of lipids moving from the planar to the source. The local triangular coordinates $u$ and $v$ are mapped across triangles such that the metric and direction of the vector are preserved. A plane cannot be mapped to an arbitrary object, like a sphere. However, for sufficiently small patches the transformation is well-defined. Local to the target point, the molecular structure corresponds closely to the source point but diverges near the patch edge. Figure S1 shows an example where the coordinate system of the planar source simulation (inset) is mapped to a coordinate system of the target fusion pore.

The lipid spacing of the source and target systems are designed to be equal at $\delta$ of target simulation, rather than at its bilayer midplane where the surface is defined. This is accomplished by an additional scaling of the $\boldsymbol{x}'$ and $\boldsymbol{y}'$ deviations from the central lipid by:

$$\Delta x'_{\text{midplane}} = (1 - \delta J_x) \Delta x'_{\text{pivotal}} \quad \text{(S5)}$$

$$\Delta y'_{\text{midplane}} = (1 - \delta J_y) \Delta y'_{\text{pivotal}}, \quad \text{(S6)}$$

where $J_x$ and $J_y$ are the curvature evaluated along the $x$ and $y$ directions, respectively. When $J$ is negative, spacings are smaller at the pivotal plane than at the midplane. To move along the pivotal plane by one distance unit requires moving along the midplane by the factor in Equation S6, $(1 - \delta J)$. Two different radial fusion pore sizes were considered to emulate a fusion pore opening / closing event: i) initial midplane radius of ~16 Å ($P_{small}$) and ii) initial midplane radius of ~34 Å ($P_{large}$). Both fusion pores were simulated ~5–10 ns using NAMD (using the same methods as in the preceding bilayer section) and then converted to dms format using `charmmgui2dms.py`.

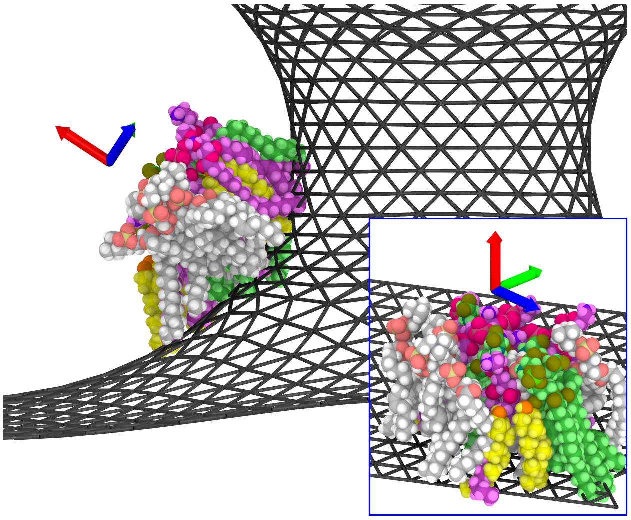

FIG. S1. An illustration of the building procedure. *Inset:* A patch of lipids from a planar source simulation is taken and inserted into a small region on the target continuum surface. The Cartesian axes are illustrated for the planar simulation with red as the $z$ axis (the average bilayer normal). *Main figure:* The target continuum mesh is shown in black. The coordinate system local to the target face is shown with red indicating the surface normal, and the $x$ and $y$ axes transformed from the planar simulation at the target point. For the patch, lipids are placed into the target's local curvilinear coordinate system. Additionally, the source lipids may be perturbed slightly (rotated and translated) to relax ring clashes and overlap.

###### *3. Anton simulation details*

The bilayer and two fusion pores were simulated using the 128-node Anton 2 supercomputer housed at the Pittsburgh Supercomputing Center (PSC), which is herein simply referred to as Anton [18]. The CHARMM all-atom lipid force field was used throughout [10]. Other than the temperature (310.15 K) and adding the semi-isotropic pressure flag, all other options were kept the same as the `basic.ark` input file supplied by the PSC. For example, the *multigrator* integration method [19], the Martyna-Tobias-Klein (MTK) [20] barostat set to 1 atm, the Nosé-Hoover thermostat [21], the M-SHAKE algorithm to constrain covalent bonds involving hydrogen [22], and a 2.5 fs timestep were used. Cutoff values were automatically determined during the Anton simulation setup and long-range electrostatics were computed using the $u$-series method [23]. We obtained a continuous ~5.6 μs for $P_{\text{small}}$, ~5.5 μs for $P_{\text{large}}$, and 2 μs for the asymmetric bilayer. The lipid redistribution (fusion pores) and $\Delta\kappa_{\text{G}}$ analyses (bilayer) were performed on these systems.

###### *4. Including carbon nanotubes as water channels in the fusion pores*

Following the long production run of the two fusion pores, both systems were modified to include two carbon nanotube (CNT) porins [24] in the bulk region at the top and bottom of the fusion pore. CNTs were built using the CHARMM-GUI *Nanomaterial Modeler* [25], targeting a radius of 10.8 Å and a length of 29.5 Å. Terminal hydrogens were replaced with hydroxyls to create a polar interface. Lipids in the fusion pore structures were removed to create cylindrical membranes holes for the embedded CNTs. Once added, CNT-lipid contacts were check to avoid clashes between neighboring acyl chains and the CNT ring structure. The systems were converted to Amber format [26, 27] using ParmEd. They were simulated for 1.57 μs ($P_{\text{small, CNT}}$) and 1.49 μs ($P_{\text{large, CNT}}$) using the Amber18 version of `pmemd.cuda` [28–30] and the CHARMM all-atom lipid force field. Constant pressure of 1 bar was maintained by a Monte Carlo barostat [31, 32]. The constant temperature of 310.15 K was maintained by Langevin dynamics with a 1 $\text{ps}^{-1}$ damping coefficient. Non-bonded interactions were switched off between 10–12 Å. Covalent bonds involving hydrogen were constrained using the SHAKE and SETTLE algorithms, and long-range electrostatics were calculated by PME.

##### C. Other bilayers to test specific hypotheses

###### *1. Bilayers to test fusion pore membrane thickness*

We observed thinned neck regions (relative to the bulk) and cholesterol exclusion in the Anton-generated fusion pore simulations. We aimed to determine whether the bulk had been thickened and the fusion pore thinned simply because of cholesterol redistribution. Four symmetric systems (compositions of: $P_{small}$'s outer leaflet, $P_{small}$'s inner leaflet, $P_{large}$'s outer leaflet, and $P_{large}$'s inner leaflet) were built and simulated 50 ns in triplicate with NAMD to obtain equilibrium leaflet areas and thicknesses. These systems were built to be relatively small (100 lipids/leaflet) so that large-scale fluctuations would not influence the thickness calculations. Then, asymmetric membranes matching the $P_{small}$ and $P_{large}$ neck compositions were built in triplicate. The "inner" leaflets had larger area/lipid on average than the "outer" leaflets, so the inner leaflet concentrations were scaled so that the areas matched. That is, the outer leaflets still contained 100 lipids, but the inner leaflets contained 100+ lipids. These asymmetric membranes were also simulated 50 ns in triplicate with NAMD as described above to obtain equilibrium areas and thicknesses used for analysis.

###### *2. Bilayers to test the thickness fluctuation method*

We introduce a method to calculate a lipid's preferred *bilayer* thickness based on thickness fluctuation spectra (see the Analysis section). Given the complexity of our plasma membrane mimetic, we also simulated POPC:chol (70:30 mol%) for a simplified system. Six independent replicas were used that contained 200 lipids per leaflet, 50 $H_2O$ per lipid, and 150 mM KCl. Following a short NAMD equilibration period, the systems were converted to Amber format using ParmEd and simulated using `pmemd.cuda` and the CHARMM all-atom lipid force field as described above. Each of the six independent replicas were simulated 2 μs, and the first 100 ns were left off for equilibration.

#### II. ANALYSIS METHODS

Throughout, the bilayer is characterized through a fit through its midplane surface. For individual lipids, curvature is analyzed at their position in the leaflet using the *neutral surface* position, $\delta$. The $\delta$ is where curvature and lateral area elasticity are independent. When measured at $\delta$, the area of leaflet is constant even as it is bent into the shape of a fusion pore. The following details how curvature, area, and thickness are determined for lipids given a description of the bilayer midplane.

##### A. Fusion pore analysis

###### *1. Fitting the all-atom fusion pore simulations*

Continuum meshes (described in Sections I A and I B 2 to construct the fusion pores) were also used to analyze the simulations. Fits to create the midplane surfaces were obtained by matching the surface to the average density of $\delta$ atoms collected from 20-nanosecond blocks of the simulation. In addition to the HC energy, an additional term was added to favor the surface midplane passing between the $\delta$s. At the same integration points used to evaluate the HC energy, the density was measured a distance $\delta$ from the midplane, for both the inner and outer leaflets. The value of $\delta$ was optimized during the simulation, but was between 12.5 and 13.1 Å for each simulation. The distance $\delta$ was modified by the local curvature to reflect the change in thickness. The recorded $\delta$ surface density was interpolated between three-dimensional grid centers, such that the gradient was continuous. For each point $\boldsymbol{r}_i$ on the surface, a term $e_i$ was added to the energy as:

$$e_i = -\frac{g_0}{n} * \lambda * \log(1 + n(\boldsymbol{r}_i)) \tag{S7}$$

where $g_0$ is the metric weight of the surface, $\lambda$ is the strength of the coupling (set to 1000 kcal/mol), and $n(\boldsymbol{r}_i)$ is the number of $\delta$ atoms in the box. The strength of the coupling was varied heuristically to yield a good fit. If the coupling is too high, the surface will crumple to fit the density. If too low, the surface will not adequately fit the density. The fit energy, containing HC and density-matching terms, was then iterated $3 \times 10^5$ times using the BFGS minimization algorithm to convergence with the Hessian reset every 100 steps. This created a series of continuum-mesh fits, one for each 20 nanosecond block of a trajectory.

###### *2. Extracting lipid curvature*

For each lipid in the trajectory, the nearest point on the corresponding fit surface was obtained. From this point, lipids are assigned to regions of the fusion pore at each point in the trajectory. A lipid's leaflet is detected depending on whether the lipid's position was above or below the midplane.

##### 3. Thickness variation with $J_{\rm mid}$ and $K_{\rm mid}$

A leaflet's thickness variation as a function of $J_{\rm mid}$ and $K_{\rm mid}$ is determined by solving for the thickness that maintains constant volume. Starting with Equation 6 in the main text, and assuming that the product of curvature and thickness is small, the leaflet's curvature a distance $\delta$ from the midplane along the surface normal is:

$$J = J_{\rm mid} - J_{\rm mid}^2 \delta + 2K_{\rm mid}\delta + \mathcal{O}[\delta^2] \tag{S8}$$

For a flat surface, the neutral surface is at $\delta_0$, and the area at this location is $A_0$. The area a distance $z$ away from $\delta$ (toward the bilayer midplane) is:

$$A(z) = A_0(1 - J(\delta)z + K(\delta)z^2) \tag{S9}$$

That is, the area decreases for positive $J$, see the blue leaflet of Figure 1 of the main text. We use $K(\delta) \approx K_{\rm mid}$ for the treatment here, sufficient for expansion of the thickness to second order in $\delta_0$. The volume is determined by integrating from the surface back to the midplane:

$$V = \int_0^\delta \mathrm{d}z \; A(z) \tag{S10}$$

$$= A_0\delta - \frac{1}{2}A_0 J_{\rm mid}\delta^2 + \frac{1}{2}A_0 J_{\rm mid}^2 \delta^3 - \frac{2}{3}A_0 K \delta^3 \tag{S11}$$

For small curvature, $\delta$ will be nearly $\delta_0$. Expanding $V$ in a series to first order in $t = \delta - \delta_0$ and solving for $V = \delta_0 A_0$ yields

$$\delta_{\rm out} = \delta_0 \left(1 + \frac{J_{\rm mid}\delta_0}{2} + \frac{2}{3}K\delta_0^2 + \mathcal{O}[\delta_0^3]\right) \tag{S12}$$

for the outer leaflet and

$$\delta_{\rm in} = \delta_0 \left(1 - \frac{J_{\rm mid}\delta_0}{2} + \frac{2}{3}K\delta_0^2 + \mathcal{O}[\delta_0^3]\right) \tag{S13}$$

for the inner leaflet. For a symmetric bilayer, the *bilayer* interior thickness within $\delta$ is the sum of that of the leaflets:

$$\delta_{\rm in} + \delta_{\rm out} = 2\delta_0 \left(1 + \frac{2}{3}K\delta_0^2 + \mathcal{O}[\delta_0^3]\right) \qquad \text{(symmetric bilayer)} \tag{S14}$$

Therefore, saddle curvature with $K < 0$ *thins* the bilayer interior relative to a planar bilayer of the same composition (Figure S2).

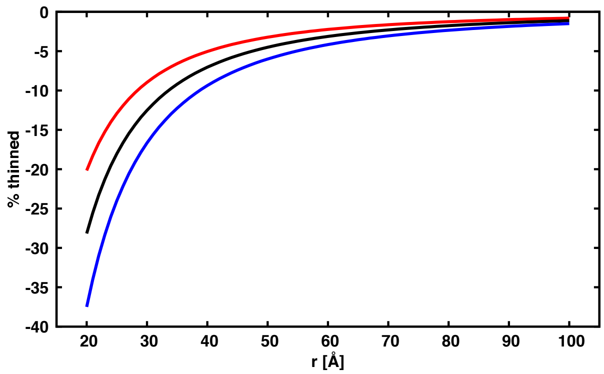


FIG. S2. Bilayer thinning as a function of radius for a fusion pore with a perfect saddle shape ($K_{\rm mid} = -1/r^2$). The theoretical thinning follows Equation S14. In this example, leaflet $\delta$ of 11 Å (red), 13 Å (black), and 15 Å (blue) were used as test values. The bilayer % thinning is calculated by: $100 \times (t_{\rm theory} - 2\delta)/t_{\rm theory}$. The thinning becomes sub-Å at $r$ ~42 Å ($\delta = 11$ Å), ~54 Å ($\delta = 13$ Å), and ~67 Å ($\delta = 15$ Å), indicating that thinning is only observable for small fusion pores.

*4. Defining the two regions using the surface fit*

With the continuum surface fit to the fusion pore (SM Sections I A and I B 2), $J$, $K$, and lipid distributions were accessible for each frame. Given the radial symmetry of the fusion pores, we reduced the geometry into two distinct regions: i) the bulk, the relatively planar and unperturbed region farthest from the pore; and ii) the neck, the interior of the pore. The two regions were determined by a hard radial cutoff from the pore center. For $P_{small}$, lipids within a radial distance $< 30$ Å were considered to be in the neck (with the other lipids in the bulk). For $P_{large}$, the radial cutoff was $<$ 60 Å. For the two regions, we then calculated the average curvatures ($\langle J \rangle$ and $\langle K \rangle$) and lipid distributions (e.g., in the main text, Figure 5).

#### B. Planar bilayer analysis

*1. Determination of the leaflet neutral surface and bilayer bending modulus*

Following the work done in Reference [33], we calculated the location of $\delta_0$ using the transverse curvature bias as a function of lipid atom. The average curvature of a planar surface is zero only if a lipid's position is measured at $\delta$. Therefore, for lipids experiencing strong negative curvature, the lipids will appear more concentrated above $\delta_0$ simply because of biased positional sampling. See Figure 1 of Reference [33] and Figure 1 of the main text.

At the leaflet $A_0$ and curvature fluctuations are uncoupled. We estimate $\delta_0$ as the height above the surface at which the experienced curvature, averaged over all lipids, is zero. For planar simulations, surfaces deviating from $\delta_0$ will display non-zero average curvature, as Fourier modes will not be sampled uniformly (that is, lipid density and curvature are coupled in violation of the $\delta_0$ condition). See Reference [33] for further details.

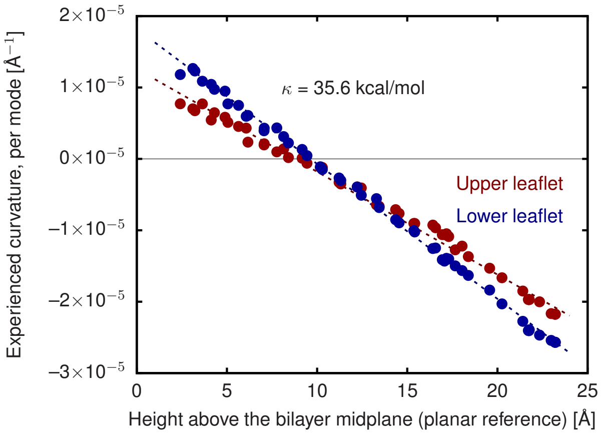


FIG. S3. Transverse curvature bias of the upper and lower leaflets from which the bending modulus is extracted.

*2. Individual lipid properties from redistribution analysis*

Lipid mechanical properties can be extracted from small fluctuations of a simulation of a planar bilayer. Fluctuations of planar simulations are conveniently described in terms of Fourier modes, allowing quantification of $q$-dependent membrane properties. To begin, the leaflet heights and total bilayer thickness $t$ are described in the Monge gauge using the height functions $h_{\rm in}$ and $h_{\rm out}$ of the inner and outer leaflet, respectively:

$$t(x, y) = [h_{\rm out}(x, y) - h_{\rm in}(x, y)] \qquad \text{(S15)}$$

and curvature $J$

$$J_{\rm in/out}(x, y) = \pm\nabla^2 h_{\rm in/out}(x, y) \qquad \text{(S16)}$$

Fluctuations of $t$ and $J$ are governed by the compressibility modulus $K_{\rm A}$ and the bending modulus $\kappa_{\rm m}$ [34]:

$$\overline{F}(J, t) = \frac{K_{\rm A}}{2}\epsilon^2 + \frac{\kappa_{\rm m}}{2}\left(((J_{\rm in} - J_0)^2 + (J_{\rm out} - J_0)^2\right), \qquad \text{(S17)}$$

where the bilayer thickness strain, $\epsilon$ is

$$\epsilon = \frac{t(x,y) - t_0}{t_0}, \tag{S18}$$

An individual lipid, whose properties $\Delta J_0$ and $\Delta t_0$ are distinct from the bulk, perturbs the energy locally [33]. We use the *leaflet* bending modulus $\kappa_{\mathrm{m}}$ instead of $\kappa$ because curvature is a property of leaflets, whereas we model thickness as a bilayer property not resolvable at the leaflet level.

$$\Delta \overline{F}_{\mathrm{p}} = \underbrace{\frac{\kappa_{\mathrm{m}}}{2}\left[J(x',y') - \Delta J_0 - J_0\right]^2 - \frac{\kappa_{\mathrm{m}}}{2}\left[J(x',y') - J_0\right]^2}_{\text{change in curvature elasticity}} + \underbrace{\frac{K_{\mathrm{A}}}{2t_0^2}\left[t(x',y') - \Delta t_0 - t_0\right]^2 - \frac{K_{\mathrm{A}}}{2t_0^2}\left[t(x',y') - t_0\right]^2}_{\text{change in thickness elasticity}}, \tag{S19}$$

Primed coordinates $\{x', y'\}$ indicate the position of the lipid. The lipid's area extent ($A_{\mathrm{p}}$) is approximated as a Dirac delta function ($\delta(\boldsymbol{r})$) with normalization

$$\int_A \mathrm{d}A \; A_{\mathrm{p}} \delta(\boldsymbol{r}) = A_{\mathrm{p}} \tag{S20}$$

Integration over the area in the vicinity of the lipid is:

$$F_{\mathrm{p}} = \int_A \mathrm{d}A \; \overline{F}_{\mathrm{p}} A_{\mathrm{p}} \delta(\{x,y\} - \{x',y'\}). \tag{S21}$$

With $h$ expressed in Fourier space,

$$h_{\mathrm{in/out}}(x,y) = A^{-1} \sum_{\boldsymbol{q}} h_{\mathrm{in/out},\boldsymbol{q}} \exp\left(\imath \boldsymbol{q} \cdot \{x,y\}\right) \tag{S22}$$

curvature is:

$$J_{\mathrm{mid}}(x,y) = -A^{-1} \sum_{\boldsymbol{q}} q^2 \frac{h_{\mathrm{out},\boldsymbol{q}} + h_{\mathrm{out},\boldsymbol{q}}}{2} \exp\left(\imath \boldsymbol{q} \cdot \{x,y\}\right) \tag{S23}$$

and bilayer thickness is

$$t(x,y) = A^{-1} \sum_{\boldsymbol{q}} \left(h_{\mathrm{out},\boldsymbol{q}} - h_{\mathrm{in},\boldsymbol{q}}\right) \exp\left(\imath \boldsymbol{q} \cdot \{x,y\}\right). \tag{S24}$$

For thermal undulations, corrections to $J_{\mathrm{mid}}$ for leaflet position are negligible; $J$ and $J_{\mathrm{mid}}$ can be used interchangeably. When squaring $J$ and $t$ in their harmonic potentials, cross-terms between modes $\boldsymbol{q}$ and $\boldsymbol{q}'$ only couple for equal $\boldsymbol{q}$ when integrating over the domain of the surface; The function $\exp\left(\imath\left(\boldsymbol{q} - \boldsymbol{q}'\right) \cdot \boldsymbol{r}\right)$ integrates to zero if $\boldsymbol{q} \neq \boldsymbol{q}'$. An individual lipid's $\Delta J_0$ is computed from

$$\langle J \rangle = Z^{-1} \mathrm{d}\{h_{\boldsymbol{q}}, x', y'\} J \exp\left(-\beta[F(J,t) + F_{\mathrm{p}}]\right) \tag{S25}$$

$$Z = \int \mathrm{d}\{h_{\boldsymbol{q}}, x', y'\} \exp\left(-\beta[F(J,t) + F_{\mathrm{p}}]\right) \tag{S26}$$

as

$$\Delta J_0 = \langle J \rangle \frac{A_p}{A} \tag{S27}$$

per mode and to first order in $\Delta J_0$ and $\Delta t_0$. Expanded to first order in $A_{\mathrm{p}}$, such that $\exp\left(-\beta[F(J,t) + F_{\mathrm{p}}]\right)$ is a Gaussian, the integrals in Equations S25 and S26 are of straightforward Gaussian type. If $\langle J \rangle$ is collected as a function of $q$ the spatial dependence of the lipid's influence can be computed [33]. Here we assume local extent and thus average $\langle J \rangle$ up to $q = 0.12$. Figure S4 shows the transverse curvature bias (the average curvature collected as a function of atom transverse position in the leaflet) for the planar simulation. Table S4 shows the difference in spontaneous curvature for each lipid, inferred from the average curvature sampled.

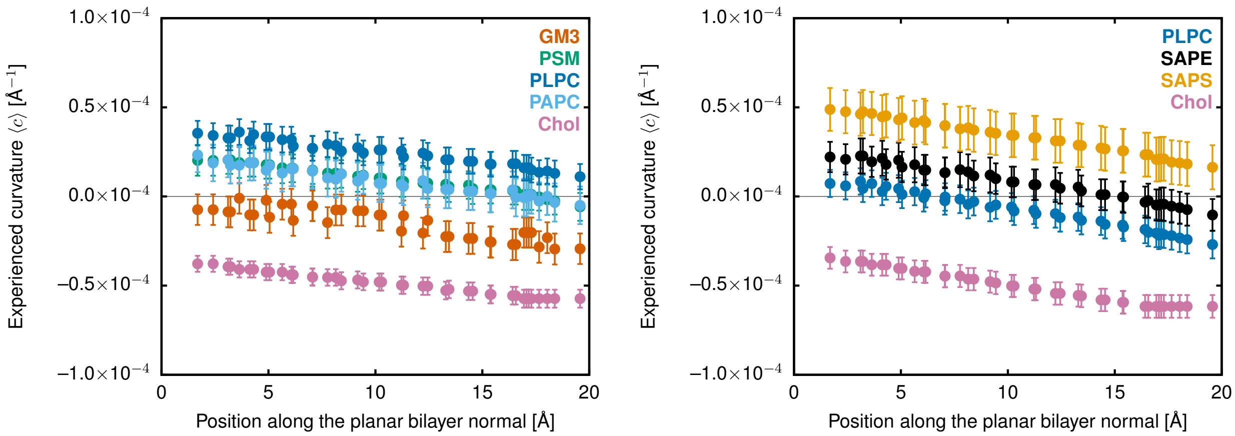


FIG. S4. Transverse curvature bias and CCR analysis of the spontaneous curvature of the upper and lower leaflets. Data are taken from the planar simulation.

| Leaflet | chol | PLPC | SAPE | SAPS | PSM | GM3 | PAPC |
|---|---|---|---|---|---|---|---|
| Inner | −0.0226 ± 0.0028 | −0.0018 ± 0.0024 | 0.0024 ± 0.0025 | 0.0100 ± 0.0036 | | | |
| Outer | −0.0217 ± 0.0022 | 0.0081 ± 0.0022 | | | 0.0038 ± 0.0032 | −0.0038 ± 0.0032 | 0.0021 ± 0.0030 |

TABLE S4. Spontaneous curvature differences ($\Delta J_0$) inferred from CCR analysis of the planar Anton simulation.

##### 3. *Thickness-coupled-redistribution (TCR) analysis*

An individual lipid's $\Delta t_0$ is computed from

$$\langle t \rangle = Z^{-1} \int \mathrm{d}\{h_{\boldsymbol{q}}, x', y'\} t(x', y') m(x', y') \exp\left(-\beta[F(J,t) + F_{\mathrm{p}}]\right) \tag{S28}$$

as

$$\langle t \rangle \approx \left( \frac{A_{\mathrm{p}}}{A} \left( 2\Delta t_0 + \frac{\kappa \Delta J_0 q^2 t_0^2}{2K_{\mathrm{A}}} \right) + \frac{2t_0}{A\beta K_{\mathrm{A}}} \right) \left( 1 + \frac{\kappa q^4 t_0^2}{4K_{\mathrm{A}}} \right)^{-1} + \mathcal{O}[\Delta J_0^2, \Delta t^2, \Delta J_0 \Delta t] \tag{S29}$$

to first order in $\Delta J_0$ and $\Delta t_0$. Here $m(x', y')$ models the increase in the density of lipids with thickness:

$$m(x', y') = 1 + \frac{t(x'y') - t_0}{t_0}. \tag{S30}$$

That is, when a lipid thickens its area decreases, assuming constant volume [35]. The strain in area corresponds with the inverse "strain" in terms of lipid density. To first order in the strain this is $m(x', y')$ above. Lipids will be sampled more frequently at thicker regions of the membrane because of the thickening. The term $\frac{2t_0}{A\beta K_{\mathrm{A}}} \left( 1 + \frac{\kappa q^4 t_0^2}{4K_{\mathrm{A}}} \right)^{-1}$ arises directly from the metric; we treat this as an artifact for which $\langle t \rangle$ must be corrected. It can either be subtracted off using the explicit formula for the term, or alternatively the average thickness deviation can be set to zero. We choose to zero the average thickness at each $q$ point. The quantity subtracted at $q$ near zero (0.0038 Å, lower; 0.0023 Å, upper) agrees well with the theoretical prediction ($\frac{2t_0}{A\beta K_{\mathrm{A}}}$ = 0.0029 Å).

The quantity $\langle t \rangle$ depends explicitly on $q$ even with local lipid extent:

$$\langle t \rangle \propto \left( 1 + \frac{\kappa q^4 t_0^2}{4K_{\mathrm{A}}} \right)^{-1}. \tag{S31}$$

The variation with $q$ is due to the different $q$ dependence between curvature ($q^4$) and thickness (no $q$ dependence). The curvature dependence of $\langle t \rangle$ ($\propto \frac{\Delta J_0 q^2 t_0^2}{4K_{\mathrm{A}}}$) is accounted for by subtracting off the contribution, using $J_0$ extracted above from CCR analysis.

The values of $\Delta t_0$ are taken by fitting Equation S29 (subtracting the term proportional to $J_0$, extracted from CCR analysis) from a histogram of $\langle t \rangle$ grouped by only two bins centered at $q = \{0.050, 0.075\}$ Å$^{-1}$ where the thickness effect

| Leaflet | chol | PLPC | SAPE | SAPS | PSM | GM3 | PAPC |
|---|---|---|---|---|---|---|---|
| Inner | 3.9 ± 1.3 | –1.3 ± 0.4 | –0.1 ± 0.4 | –1.3 ± 0.4 | | | |
| Outer | 1.8 ± 0.8 | –0.1 ± 0.9 | | | 0.2 ± 0.8 | –2.5 ± 1.2 | –1.5 ± 0.7 |

TABLE S5. Bilayer thickness-preference differences ($\Delta t_0$, Å) inferred from TCR analysis of the planar Anton simulation.

| Leaflet | chol | PLPC | SAPE | SAPS | PSM | GM3 | PAPC |
|---|---|---|---|---|---|---|---|
| Inner | –10.4 ± 3.5 | 3.4 ± 1.0 | 0.2 ± 1.0 | 3.5 ± 1.1 | | | |
| Outer | –4.7 ± 2.2 | 0.1 ± 2.3 | | | –0.6 ± 2.2 | 6.8 ± 3.2 | 4.0 ± 1.8 |

TABLE S6. Predicted changes in the Gaussian curvature modulus ($\Delta\kappa_G$, kcal/mol) on the basis of bilayer thickness elasticity.

is dominant, and where molecular-scale effects are least significant. That is, while the lower $q$ points have the largest stochastic error, they are the most clearly valid, in terms of continuum mechanics; the wavelength of the mode is larger than the thickness of the bilayer. Note that the complete simulation observations are inconsistent with the stochastic error in the measurement of $\langle t \rangle$ (the model does not fully explain the data). This strongly suggests there is $q$-dependence to the thickness, for example, by coupling to the size of small, thick domains that form in the outer leaflet, or by molecular degrees of freedom not accounted for by the simple theory.

Energetic terms proportional to $K$ are not treated explicitly in this model. Rather, $K$ is used through Equations 5 or 6 to compute a lipid's curvature at its $\delta$. which is treated in our model through the *leaflet* $J$ dependence on $K$. For our initial model we assume that $\kappa_{\mathrm{G,m}}$ is not a localized property, similar to our treatment of $\kappa_{\mathrm{m}}$. That is, $\kappa_{\mathrm{m}}$ and $\kappa_{\mathrm{G,m}}$ might vary between phases like $\mathrm{L_o}$ and $\mathrm{L_d}$, but that a single lipid doesn't have a localized stiffness.

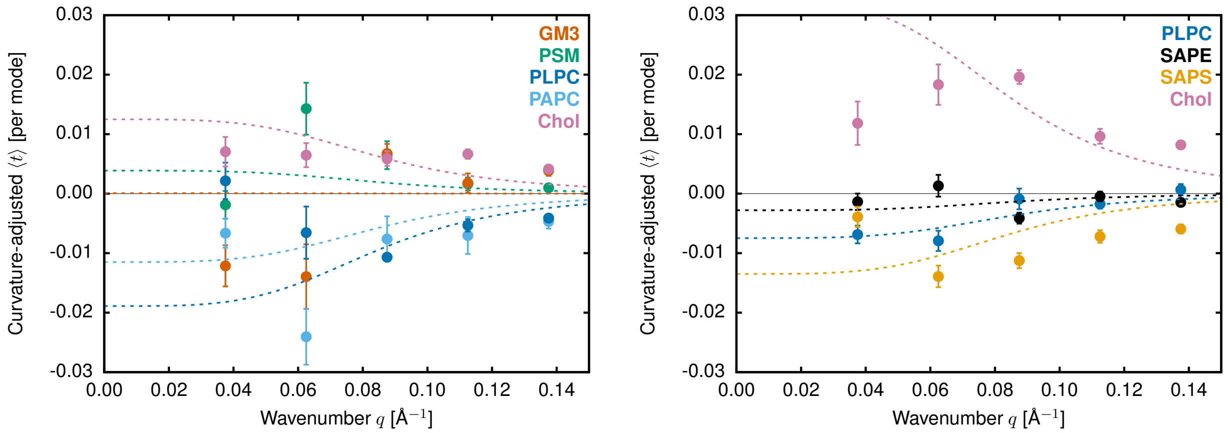


FIG. S5. $q$-dependent TCR analysis of the preferred thickness of the upper (left) and lower (right) leaflets. Solid points are taken from the planar simulation. Dashed lines are best-fit.

###### 4. *Chol's bilayer thickness preference in POPC*

The TCR analysis for the complex, asymmetric planar bilayer showed that the method has difficulty describing thickness fluctuations at low $q$, which are slowly fluctuating thickness modes (Figure S5; see Figure 2 for a schematic of thickness modes). To demonstrate that the TCR method is reasonable, particularly at higher $q$, we used a simplified, symmetric system of POPC:chol at a 70:30 mol ratio. The error bars are still relatively large at low $q$, but the theoretical fit is much better. Longer timescale simulations, even for this simple lipid mixture, are necessary to have well-fit low-$q$ data.

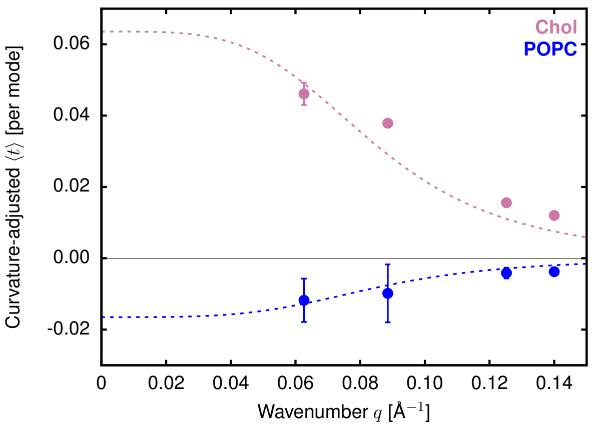


FIG. S6. $q$-dependent TCR analysis of the preferred thickness for POPC and chol in a POPC:chol (70:30) bilayer. Solid points are taken from the planar simulations. Dashed lines are best-fit.

##### 5. *The area compressibility modulus, $K_{\rm A}$*

The area compressibility modulus is computed from a planar bilayer's projected area fluctuations using the model energy:

$$F_{\rm A} = \frac{K_{\rm A}}{2} A_0 \epsilon_{\rm A}^2 \tag{S32}$$

$$= \frac{K_{\rm A}}{2A_0}(A - A_0)^2 \tag{S33}$$

where $\epsilon_{\rm A} = \frac{A-A_0}{A_0}$, and the total energy has been integrated over the entire surface. The Boltzmann distribution of states, $\exp(-\beta F_{\rm A})$, yields a Gaussian distribution with:

$$\langle (A - A_0)^2 \rangle = \frac{A_0}{K_{\rm A}\beta} \tag{S34}$$

such that

$$K_{\rm A} = \frac{k_{\rm B} T A_0}{\langle (A - A_0)^2 \rangle} \tag{S35}$$

where $\langle (A - A_0)^2 \rangle$ is the standard deviation ($\sigma_A$) of $A$. From $\sigma_A$ and $A_0$, $K_{\rm A}$ is 0.62 $\pm$ 0.03 kcal/mol/Å$^{-2}$ (431 $\pm$ 21 mN/m).

##### 6. *Bilayer thickness with fusion pore neck composition*

From the simulations setup in Section I C 1, we obtained planar $\delta_{0,\rm b}$ of 25.3 $\pm$ 0.1 Å and 26.5 $\pm$ 0.1 Å that match the pore compositions of P$_{\rm small}$ and P$_{\rm large}$, respectively. The bulk $\delta_{\rm b}$ of P$_{\rm small}$ and P$_{\rm large}$ (far from the pores' necks) are 26.7 Å and 26.3 Å, respectively (Figure 6). However, the $\delta_{0,\rm b}$ from the planar bilayer simulations do modify the strain implied by the thinning shown in Figure 6 because the thinned leaflet should be compared to the thickness of a planar simulation with the same composition. The observed thinning decreases slightly from 39% to 36% for the small pore and is nearly unchanged for the larger.

#### C. Monte Carlo (literature Helfrich) model

Monte Carlo (MC) numerical simulations yielded the probability of finding a lipid $i$ in region $r$ ($p_{\rm i,r}$) with literature values of $\kappa$, $K_{\rm A}$, $J_0$, $\widetilde{A}$, $\Delta\kappa_{\rm G}$, the regional $J$ and $K$ found from the all-atom simulations, and the number of lipids in the simulations. The bending ($F_{\rm bend}$) and stretching ($F_{\rm stretch}$) contributions to the total free energy ($F_{\rm total}$) are:

$$F_{\rm bend} = \sum_{i=1}^{n_{\rm lipids}} \sum_{r=1}^{n_{\rm regions}} \frac{1}{2}\left(\frac{1}{2}\kappa_{\rm i}\right)\widetilde{A}_{\rm i} n_{\rm i,r}(J_{0,\rm i}^2 - 2J_{0,\rm i}J_{\rm r}) + \widetilde{A}_{\rm i} n_{\rm i,r}\Delta\kappa_{\rm G,i}K_{\rm r} \tag{S36}$$

$$F_{\text{stretch}} = \sum_{r=1}^{n_{\text{regions}}} \frac{1}{2}\left(\frac{1}{2}K_{\text{A,i}}\right)A_{0,\text{r}}\left(\frac{A_{\text{r}} - A_{0,\text{r}}}{A_{0,\text{r}}}\right)^2 \tag{S37}$$

$$F_{\text{total}} = F_{\text{bend}} + F_{\text{stretch}}, \tag{S38}$$

where the summation is over the curvature regions ($n_{\text{regions}}$; e.g., the bulk and neck) and the number of lipids ($n_{\text{lipids}}$). $\widetilde{A}_{\text{i}}$ is lipid $i$'s lateral area, $J_{0,\text{i}}$ is the $i$'s intrinsic curvature, $J_{\text{r}}$ is the measured average curvature of region $r$ from the MD simulation, $n_{\text{i,r}}$ is the number of $i$'s lipid type in region $r$ (e.g., number of chol in the neck), $\kappa_{\text{i}}$ is the *bilayer* bending modulus of lipid $i$ ($\kappa_{\text{i}} = 2\kappa_{\text{m,i}}$ where $\kappa_{\text{m,i}}$ is the leaflet bending modulus), $\Delta\kappa_{\text{G,i}}$ is the lipid's Gaussian modulus *bilayer* thickness component, and $K_{\text{r}}$ is the measured average $K_{\text{mid}}$ of region $r$ from the MD simulation.

Equation S37 is the stretching contribution to the free energy and includes the compression modulus ($K_{\text{A,i}}$ (see the preceding subsection); multiplied by 1/2 to obtain a leaflet quantity), $A_{0,\text{r}}$ is the desired equilibrium area of lipids in region $r$:

$$A_{0,\text{r}} = \sum_{i=1}^{n_{\text{lipids}}} \widetilde{A}_{\text{i}} n_{\text{i,r}}, \tag{S39}$$

and $A_{\text{r}}$ is the region's inflexible neutral surface area defined by analyses of the MD trajectories' midplanes and Equation 2. A difference between $A_{0,\text{r}}$ and $A_{\text{r}}$ yields an area strain that is penalized, thereby restricting the number of lipids in each region. The total free energy ($F_{\text{total}}$) is the summation of the bending and stretching contributions.

The MC was setup with $J$ and $K$ calculated for the inner or outer leaflet of $P_{\text{large}}$ or $P_{\text{small}}$. First, lipids were randomly assigned to a region and the energy was evaluated based on Equation S38 and the parameters listed in Table S7. A random lipid $i$ was selected to move into a random region, $r$. The region was determined by a random selection based on area (i.e., it is twice more likely to move into a region with twice the area). The move was accepted/rejected by the common Metropolis criterion (the move is accepted if a new random number is less than $p = \exp(-(F_{\text{new}} - F_{\text{old}})/k_{\text{B}}T))$. Note that the entropy contribution to the free energy is naturally included in the MC sampling. This procedure was repeated for $10^6$ steps (leaving off the first $10^5$ steps as equilibration) for 10 replicas, which converged each region's lipid concentrations to a standard error of $<$0.1%.

| Lipid | $J_0$ [Å$^{-1}$] | $\widetilde{A}$ [Å$^2$] | $\kappa$ [kcal/mol] | $K_{\text{A}}$ [kcal/mol/Å$^2$] |
|---|---|---|---|---|
| GM3 | 0.00926 | 55.1 | 35.6 | 0.62 |
| chol | –0.03300 | 40.0 | 35.6 | 0.62 |
| PAPC | –0.00813 | 70.8 | 35.6 | 0.62 |
| PLPC | –0.00318 | 65.9 | 35.6 | 0.62 |
| PSM | 0.00926 | 55.1 | 35.6 | 0.62 |
| SAPE | –0.02500 | 69.3 | 35.6 | 0.62 |
| SAPS | –0.02000 | 69.3 | 35.6 | 0.62 |

TABLE S7. Parameter values derived from the literature ($J_0$, $\widetilde{A}$, and $K_{\text{A}}$) used to inform the MC model. The $\kappa$ for all lipids is assumed to be 20 kcal/mol. Note that: i) GM3 and PSM $J_0$, $\widetilde{A}$, and $K_{\text{A}}$ values are set to the PSM values taken directly from Reference [36]; ii) chol's $J_0$ is based on Reference [37, 38] and the $\widetilde{A}$ and $K_{\text{A}}$ are guess values; iii) PAPC and SAPE are set to be $J_0$, $\widetilde{A}$, and $K_{\text{A}}$ values are set to those of PDPC and SDPE, respectively [36]; iv) SAPS is set with a less negative $J_0$ than SAPE based on evidence from Reference [33] (the $\widetilde{A}$ and $K_{\text{A}}$ are set to the same as SAPE). The value of $\Delta\overline{\kappa}_{\text{chol}}$ is discussed in the main text and tested by the MC model.

The MC fits to MD were improved by adding $\Delta\kappa_{\text{G,chol}}$ to the energy function, however, there were still discrepancies chol concentrations (Figure 7). Using $\Delta\kappa_{\text{G,chol}}$ minus one standard error (–13.9 kcal/mol) further improves the MC and MD chol comparison and does not adversely affect the other values (Figure S7).

#### III. WATER FLOW AND CHOLESTEROL FLIP-FLOP IN THE FUSION PORE SIMULATIONS

To fully relax the shape of the membrane, water must flow between compartments of the simulation and lipids must change leaflet. While the shape of the pores reported here are not fully relaxed, water flow and cholesterol flipping indicate

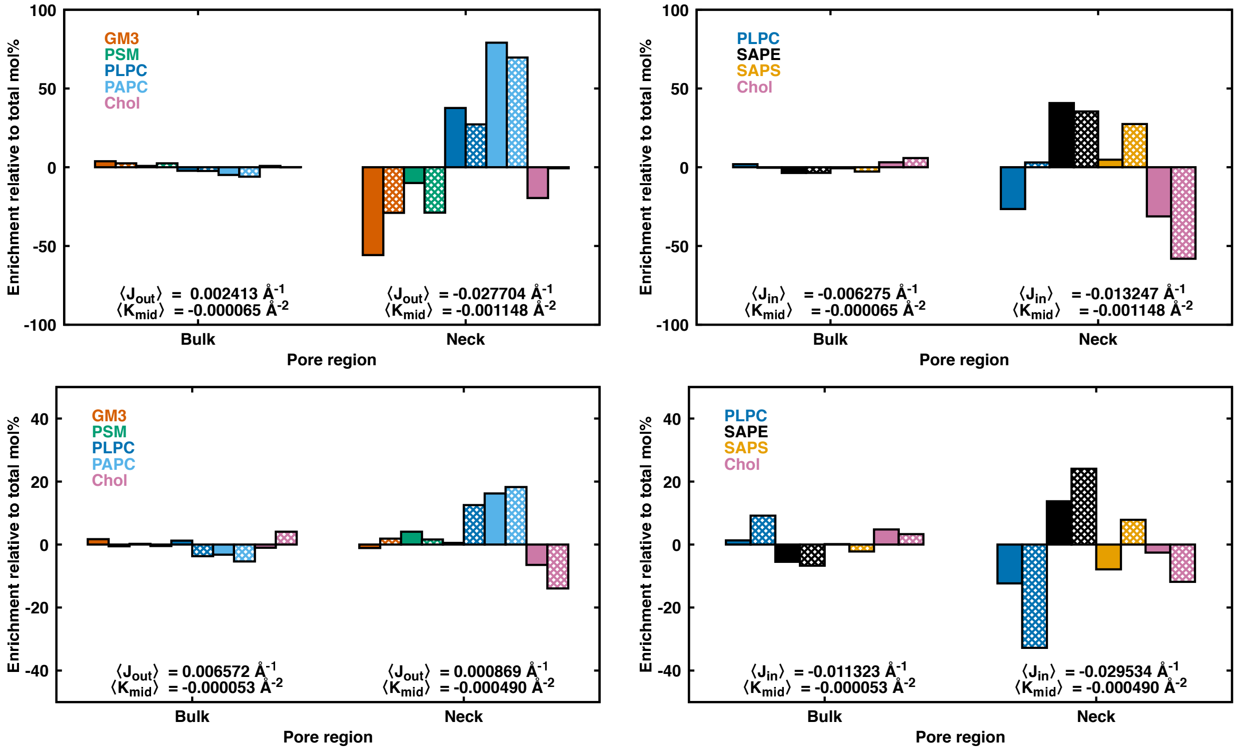


FIG. S7. Enrichment per region relative to the total mol% of a lipid species including $\Delta\kappa_{\mathrm{G,chol}}$ minus one standard error (enrichment $= 100\times(\phi_{\mathrm{species,observed}} - \phi_{\mathrm{species,total}})/\phi_{\mathrm{species,total}}$). Data for the small pore's outer (left) and inner (right) leaflets on the top row, respectively. Data for the large pore's outer (left) and inner (right) leaflets on the bottom row, respectively. See Table S1 for $\phi_{\mathrm{species,total}}$ values. Direct observations from MD are solid bars and MC predictions are hatched.

the direction of the force on the pore diameter due to elasticity. Flip-flop events recorded in the small (left) and large (right) pores are shown in Figure S8 (pre-CNTS, top; with-CNTs, bottom). Quantification of water flow through the CNT is plotted in Figure S9. In the smaller pore, water flows from the interior compartment to the outside. In the larger pore, the flow is opposite. This suggests that the equilibrium shape of the pore is in between the small and large pores. However, the flipping of cholesterol implies not only differential tension between leaflets but also chemical potential; it is an ambiguous indicator [12, 39].

Initially, $\mathrm{P_{large}}$ has 1319 lipids in its inner leaflet and 1895 lipids in its outer. From the fit to the midplane, the total area is $A_{\mathrm{mid}} = 7.96\times10^4$ Å$^2$, while average curvature $\langle J_{\mathrm{mid}}\rangle = -0.0095$ Å$^{-1}$ and average Gaussian curvature $\langle K_{\mathrm{mid}}\rangle = -0.000157$ Å$^{-2}$. $\mathrm{P_{small}}$ had 1353 lipids in its inner leaflet and 1628 lipids in its outer. From the fit to the midplane, the total area is $A_{\mathrm{mid}} = 7.48\times10^4$ Å$^2$, while average curvature $\langle J_{\mathrm{mid}}\rangle = -0.0032$ Å$^{-1}$ and average Gaussian curvature $\langle K_{\mathrm{mid}}\rangle = -0.000167$ Å$^{-2}$. Applying Equation 2 to compute the ratio $\frac{A_{0,\mathrm{in}}}{A_{0,\mathrm{out}}}$ yields 0.78 for the large pore, and 0.91 for the small pore. That is, regardless of the area of the outer pore lipids (which is reduced from compositionally pure values by the ordering effect of cholesterol on saturated tails), $\mathrm{P_{small}}$ has a smaller differential area between the inner and outer. For $\mathrm{P_{small}}$ to expand, lipids must flip from the outer leaflet to the inner. Expansion of $\mathrm{P_{small}}$ is consistent with the seven net flips of cholesterol to the outer leaflet (before CNT addition) and twelve in the 1.5 μs after CNT addition, out of sixteen total flips. In contrast, while the outer pore also has seven net flips prior to CNT addition, in the 1.5 μs following CNT addition there are zero net flips. For a strained leaflet differential where one leaflet has $\frac{1}{2}\Delta n$ lipids too many, and the other leaflet has $\frac{1}{2}\Delta n$ too few,

$$F_{\mathrm{A}} = \frac{1}{2}K_{\mathrm{A,m}}A\left(\frac{+\frac{1}{2}\Delta n\tilde{A}}{A}\right)^2 + \frac{1}{2}K_{\mathrm{A,m}}A\left(\frac{-\frac{1}{2}\Delta n\tilde{A}}{A}\right)^2,$$

$$= K_{\mathrm{A,m}}\frac{(\tilde{A}\Delta n)^2}{4A} \qquad \text{(S40)}$$

where $K_{\mathrm{A,m}}$ is the single-leaflet compressibility modulus, $\tilde{A}$ is the area per lipid, $A$ is the total area, and the energy $F_{\mathrm{A}}$ is the sum of the two leaflets' model compressibility energy. For $\mathrm{P_{small}}$ after CNT addition, twelve of sixteen flips are from the inner to outer leaflet ($p = 0.75$). Given $K_{\mathrm{A,m}}$, $\tilde{A}$, and $A$, this corresponds to $\Delta n = 10$, that is, roughly equivalent to the number of flips observed. With uncertainty in the statistics of flipping, $K_A$, as well as the unknown chemical potential for cholesterol between the leaflets, a detailed comparison is not in the scope of this work.

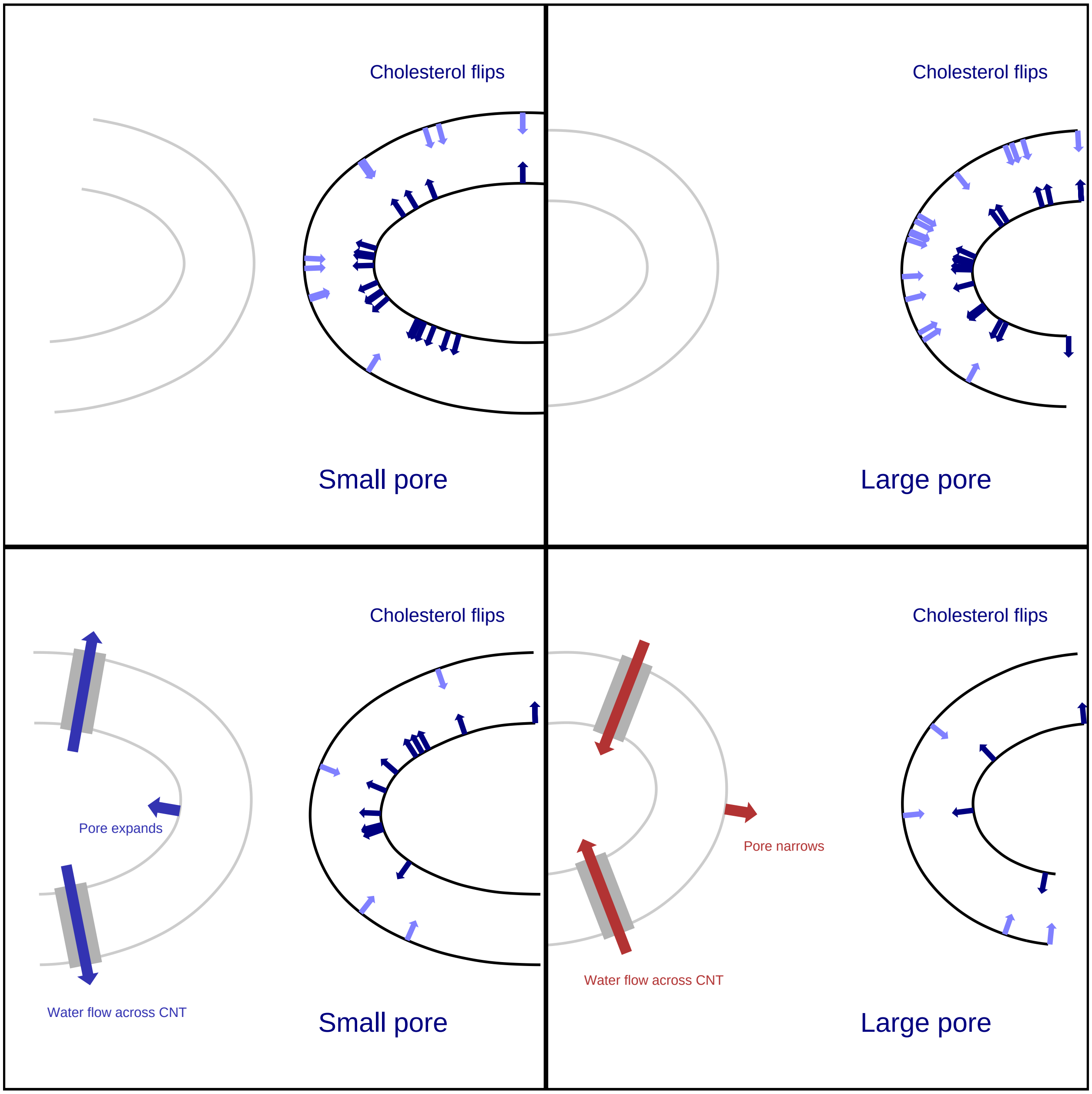


FIG. S8. A recording of flip-flop events referenced to the pore geometry where they occurred. Black arrows are flops from the inner leaflet to the outer, while blue are from the outside in. Simulations in the top row were run before addition of CNTs, for over five microseconds. Simulations in the bottom row, which include CNTs, were run for ~1.5 µs.

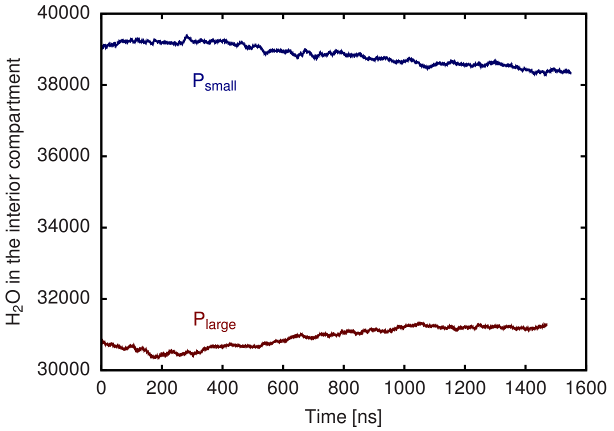


FIG. S9. $H_2O$ flow through the CNTs of the small (blue) and large (red) pores. The number of $H_2O$ in the interior compartment is plotted. For $P_{small}$, there were 39041 $H_2O$ initially in the compartment and 38362 at the final step. For $P_{large}$, there were 30828 $H_2O$ initially in the compartment and 31263 at the final step.

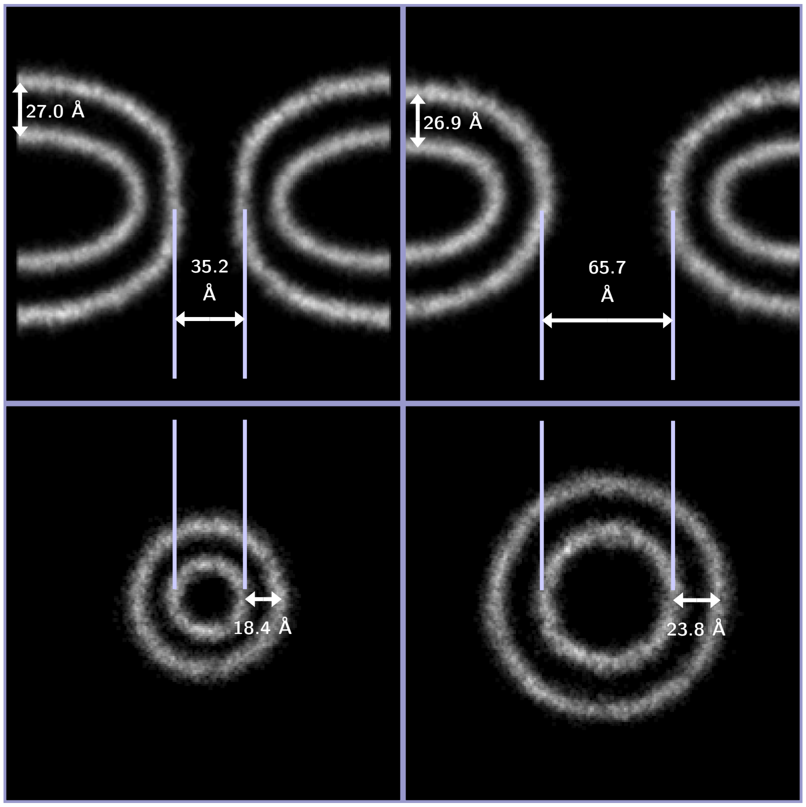


FIG. S10. The density of $\delta$ atoms for a 5-Å-wide slice, for simulations with CNTs included to relieve hydrostatic pressure. Left column: the smaller pore. Right column: the larger pore. The top row is a 5-Å-wide slice in $xz$ slice through $y = 0$. The bottom row is a 5-Å-wide slice in $xy$ through the pore at $z = 0$.

[1] F. Cirak, M. Ortiz, and P. Schröder, International Journal for Numerical Methods in Engineering **47**, 2039 (2000).
[2] F. Feng and W. S. Klug, Journal of Computational Physics **220**, 394 (2006).
[3] J. Stam, in *Proceedings of the 25th Annual Conference on Computer Graphics and Interactive Techniques, SIGGRAPH 1998* (1998) pp. 395–404.
[4] S. Jo, T. Kim, and W. Im, PLoS ONE **2**, e880 (2007).
[5] S. Jo, T. Kim, V. G. Iyer, and W. Im, Journal of Computational Chemistry **29**, 1859 (2008).
[6] S. Jo, J. B. Lim, J. B. Klauda, and W. Im, Biophysical Journal **97**, 50 (2009).
[7] J. Lee, X. Cheng, J. M. Swails, M. S. Yeom, P. K. Eastman, J. A. Lemkul, S. Wei, J. Buckner, J. C. Jeong, Y. Qi, S. Jo, V. S. Pande, D. A. Case, C. L. Brooks, A. D. MacKerell, J. B. Klauda, and W. Im, Journal of Chemical Theory and Computation **12**, 405 (2016).
[8] J. C. Phillips, R. Braun, W. Wang, J. Gumbart, E. Tajkhorshid, E. Villa, C. Chipot, R. D. Skeel, L. Kalé, and K. Schulten, "Scalable molecular dynamics with NAMD," (2005).
[9] J. C. Phillips, D. J. Hardy, J. D. Maia, J. E. Stone, J. V. Ribeiro, R. C. Bernardi, R. Buch, G. Fiorin, J. Hénin, W. Jiang, R. McGreevy, M. C. Melo, B. K. Radak, R. D. Skeel, A. Singharoy, Y. Wang, B. Roux, A. Aksimentiev, Z. Luthey-Schulten, L. V. Kalé, K. Schulten, C. Chipot, and E. Tajkhorshid, Journal of Chemical Physics **153**, 44130 (2020).
[10] J. B. Klauda, R. M. Venable, J. A. Freites, J. W. O'Connor, D. J. Tobias, C. Mondragon-Ramirez, I. Vorobyov, A. D. MacKerell, and R. W. Pastor, Journal of Physical Chemistry B **114**, 7830 (2010).
[11] S. Park, A. H. Beaven, J. B. Klauda, and W. Im, Journal of Chemical Theory and Computation **11**, 3466 (2015).
[12] A. Hossein and M. Deserno, Biophysical Journal **118**, 624 (2020).
[13] S. E. Feller, Y. Zhang, R. W. Pastor, and B. R. Brooks, The Journal of Chemical Physics **103**, 4613 (1995).
[14] J. P. Ryckaert, G. Ciccotti, and H. J. Berendsen, Journal of Computational Physics **23**, 327 (1977).
[15] S. Miyamoto and P. A. Kollman, Journal of Computational Chemistry **13**, 952 (1992).
[16] T. Darden, D. York, and L. Pedersen, The Journal of Chemical Physics **98**, 10089 (1993).
[17] U. Essmann, L. Perera, M. L. Berkowitz, T. Darden, H. Lee, and L. G. Pedersen, The Journal of Chemical Physics **103**, 8577 (1995).
[18] D. E. Shaw, J. P. Grossman, J. A. Bank, B. Batson, J. A. Butts, J. C. Chao, M. M. Deneroff, R. O. Dror, A. Even, C. H. Fenton, A. Forte, J. Gagliardo, G. Gill, B. Greskamp, C. R. Ho, D. J. Ierardi, L. Iserovich, J. S. Kuskin, R. H. Larson, T. Layman, L. S. Lee, A. K. Lerer, C. Li, D. Killebrew, K. M. Mackenzie, S. Y. H. Mok, M. A. Moraes, R. Mueller, L. J. Nociolo, J. L. Peticolas, T. Quan, D. Ramot, J. K. Salmon, D. P. Scarpazza, U. Ben Schafer, N. Siddique, C. W. Snyder, J. Spengler, P. T. P. Tang, M. Theobald, H. Toma, B. Towles, B. Vitale, S. C. Wang, and C. Young, in *International Conference for High Performance Computing, Networking, Storage and Analysis, SC*, Vol. 2015-Janua (2014) pp. 41–53.
[19] R. A. Lippert, C. Predescu, D. J. Ierardi, K. M. Mackenzie, M. P. Eastwood, R. O. Dror, and D. E. Shaw, Journal of Chemical Physics **139**, 164106 (2013).
[20] G. J. Martyna, D. J. Tobias, and M. L. Klein, The Journal of Chemical Physics **101**, 4177 (1994).
[21] S. Nosé, The Journal of Chemical Physics **81**, 511 (1984).
[22] V. Kräutler, W. F. Van Gunsteren, and P. H. Hünenberger, Journal of Computational Chemistry **22**, 501 (2001).
[23] C. Predescu, A. K. Lerer, R. A. Lippert, B. Towles, J. P. Grossman, R. M. Dirks, and D. E. Shaw, Journal of Chemical Physics **152**, 84113 (2020), arXiv:1911.01377.
[24] M. Vögele, J. Köfinger, and G. Hummer, Faraday Discussions **209**, 341 (2018).
[25] Y. K. Choi, N. R. Kern, S. Kim, K. Kanhaiya, Y. Afshar, S. H. Jeon, S. Jo, B. R. Brooks, J. Lee, E. B. Tadmor, H. Heinz, and W. Im, Journal of Chemical Theory and Computation **18**, 479 (2022).
[26] D. A. Case, T. E. Cheatham, T. Darden, H. Gohlke, R. Luo, K. M. Merz, A. Onufriev, C. Simmerling, B. Wang, and R. J. Woods, Journal of Computational Chemistry **26**, 1668 (2005).
[27] D.A. Case, I.Y. Ben-Shalom, S.R. Brozell, D.S. Cerutti, T.E. Cheatham, V. C. III, T.A. Darden, R.E. Duke, D. Ghoreishi, M.K. Gilson, H. Gohlke, A.W. Goetz, D. Greene, R Harris, N. Homeyer, S. Izadi, A. Kovalenko, T. Kurtzman, T.S. Lee, S. LeGrand, P. Li, C. Lin, J. Liu, T. Luchko, R. Luo, D.J. Mermelstein, K.M. Merz, Y. Miao, G. Monard, C. Nguyen, H. Nguyen, I. Omelyan, A. Onufriev, F. Pan, R. Qi, D.R. Roe, A. Roitberg, C. Sagui, S. Schott-Verdugo, J. Shen, C.L. Simmerling, J. Smith, R. Salomon-Ferrer, J. Swails, R.C. Walker, J. Wang, H. Wei, R.M. Wolf, X. Wu, L. Xiao, D.M. York, and P.A. Kollman, University of California, San Francisco (2018).
[28] A. W. Götz, M. J. Williamson, D. Xu, D. Poole, S. Le Grand, and R. C. Walker, Journal of Chemical Theory and Computation **8**, 1542 (2012).
[29] R. Salomon-Ferrer, A. W. Götz, D. Poole, S. Le Grand, and R. C. Walker, Journal of Chemical Theory and Computation **9**, 3878 (2013).
[30] S. Le Grand, A. W. Götz, and R. C. Walker, Computer Physics Communications **184**, 374 (2013).
[31] K.-H. Chow and D. M. Ferguson, Computer Physics Communications **91**, 283 (1995).
[32] J. Åqvist, P. Wennerström, M. Nervall, S. Bjelic, and B. O. Brandsdal, Chemical Physics Letters **384**, 288 (2004).
[33] K. C. Sapp, A. H. Beaven, and A. J. Sodt, Physical Review E **103**, 042413 (2021).
[34] H. W. Huang, Biophysical Journal **50**, 1061 (1986).
[35] M. M. Terzi, M. Deserno, and J. F. Nagle, Soft Matter **15**, 9085 (2019).
[36] R. M. Venable, F. L. Brown, and R. W. Pastor, Chemistry and Physics of Lipids **192**, 60 (2015).
[37] Z. Chen and R. P. Rand, Biophysical Journal **73**, 267 (1997).
[38] A. J. Sodt, R. M. Venable, E. Lyman, and R. W. Pastor, Physical Review Letters **117**, 138104 (2016).

[39] D. W. Allender, A. J. Sodt, and M. Schick, Biophysical Journal **116**, 2356 (2019).
